# Self-organisation of mortal filaments: the role of FtsZ treadmilling in bacterial division ring formation

**DOI:** 10.1101/2023.05.08.539808

**Authors:** Christian Vanhille-Campos, Kevin D. Whitley, Philipp Radler, Martin Loose, Séamus Holden, Anđela Šarić

## Abstract

Protein filaments in the cell commonly treadmill – they grow on one end while shrinking on the other, driven by energy consumption. Treadmilling filaments appear to be moving, even though individual proteins remain static. Here, we investigate the role of treadmilling, implemented as dynamic turnover, in the collective filament self-organisation. On the example of the bacterial FtsZ protein, a highly conserved tubulin homologue, we show, in computer simulations and *in vitro* experiments, that treadmilling drives filament nematic ordering by dissolving misaligned filaments. We demonstrate that ordering via local dissolution allows the system to quickly respond to chemical and geometrical biases in the cell, and is necessary for the formation of the FtsZ ring required for bacterial cell division in living *Bacillus subtilis* cells. We finally use simulations to quantitatively explain the characteristic dynamics of FtsZ ring formation *in vivo*. Beyond FtsZ and other cytoskeletal filaments, our study identifies a novel mechanism for nematic ordering via constant birth and death of energy-consuming filaments.

---

Cytoskeletal filaments are active cellular elements that continuously grow and shrink through addition and removal of subunits. One prominent example of this activity is treadmilling – a process by which protein filaments grow on one end, while the other end shrinks at equal rate. This dynamic behaviour is driven by nucleotide hydrolysis, and results in the filaments seemingly moving, despite the individual filament monomers remaining in one place [1]. Many essential cytoskeletal filaments, such as actin, microtubules and FtsZ, exhibit this important feature [2–6].

From a physics point of view, treadmilling has been often modelled as self-propulsion, where the filament centre of mass moves directionally with a certain velocity to mimic the directional filament growth [7–9]. While this approach is appropriate to describe single filament dynamics, when it comes to the assembly into higher order dynamic structures, self-propelled models might not be suitable. In particular, such models fail to capture the characteristic constant turnover of components in treadmilling systems, and by introducing propelling forces they may overestimate the forces generated by filament polymerisation on obstacles, which have been shown to be small or negligible [10] and instead likely operate via a ratchet-like interplay between polymerisation and thermal fluctuations [11]. The self-organisation mechanisms of treadmilling polymers may hence be very different from those of self-propelled filaments and remain unexplored.

Here we develop the first computational model for collective behaviour of treadmilling filaments, accounting for the kinetics of nucleation, growth and shrinkage, to study their collective dynamics. We particularly focus on investigating self-organisation of FtsZ filaments, a highly conserved tubulin homolog [12], widely present in bacteria, and one of the best characterised treadmilling systems in the literature [13–16]. Our model allows us to explore the interplay between polymerisation dynamics and collective filament organisation at the cellular scale, and to directly compare to *in vitro* and *in vivo* experiments. Treadmilling FtsZ filaments self-organise into a dynamic ring [17] in the middle of bacterial cells (the “Z-ring”) that orchestrates bacterial cell division [18–20]. Treadmilling has been shown to be essential for cell division of *E. coli, B. subtilis, S*.*aureus* and *S. pneumoniae* [15, 16, 21, 22] and in *B. subtilis* it has been shown to be required for the formation of a coherent Z-ring [23]. However, how treadmilling contributes to the self-organization of individual filaments into a large scale cytoskeletal structure remains unclear.

Here, we first show that our model correctly reproduces single-filament treadmilling dynamics collected in new *in vitro* measurements [13], and proposed in previous theoretical kinetic studies [24]. We then demonstrate that treadmilling polymers collectively align due to shrinkage and “death” of misaligned filaments, yielding quantitative matching with high-speed AFM imaging of reconstituted FtsZ filaments. Finally, we find that such a system forms tight ordered dynamic rings when under external biases present in bacterial division. The model quantitatively explains the time-dependent *in vivo* dynamics of Z-ring condensation and maturation [23, 25] in *B. subtilis*. These results highlight a new model for ordering of cytoskeletal filaments through dynamic growing and shrinking, which is responsible for the formation of the bacterial division ring, and potentially present in a plethora of other cytoskeletal processes across the tree of life.

## Model description

We model treadmilling filaments as coarse-grained polymers made of spherical beads with diameter *σ* (the simulation unit of length) on a two-dimensional plane with periodic boundary conditions. Individual beads correspond to distinct monomers, with *σ* = 5 nm as known for FtsZ [19, 26, 27]. Filaments are formed by connecting monomers via harmonic springs and angle potentials to capture filament stiffness (see SI for more details on the model). We evolve our system in Molecular Dynamics simulations and impose filament nucleation, growth and shrinkage kinetics by dynamically creating and deleting monomers in the system (Figure 1a and Supplementary Movie 1 – see SI for more details on the simulation setup).

**FIG. 1.**
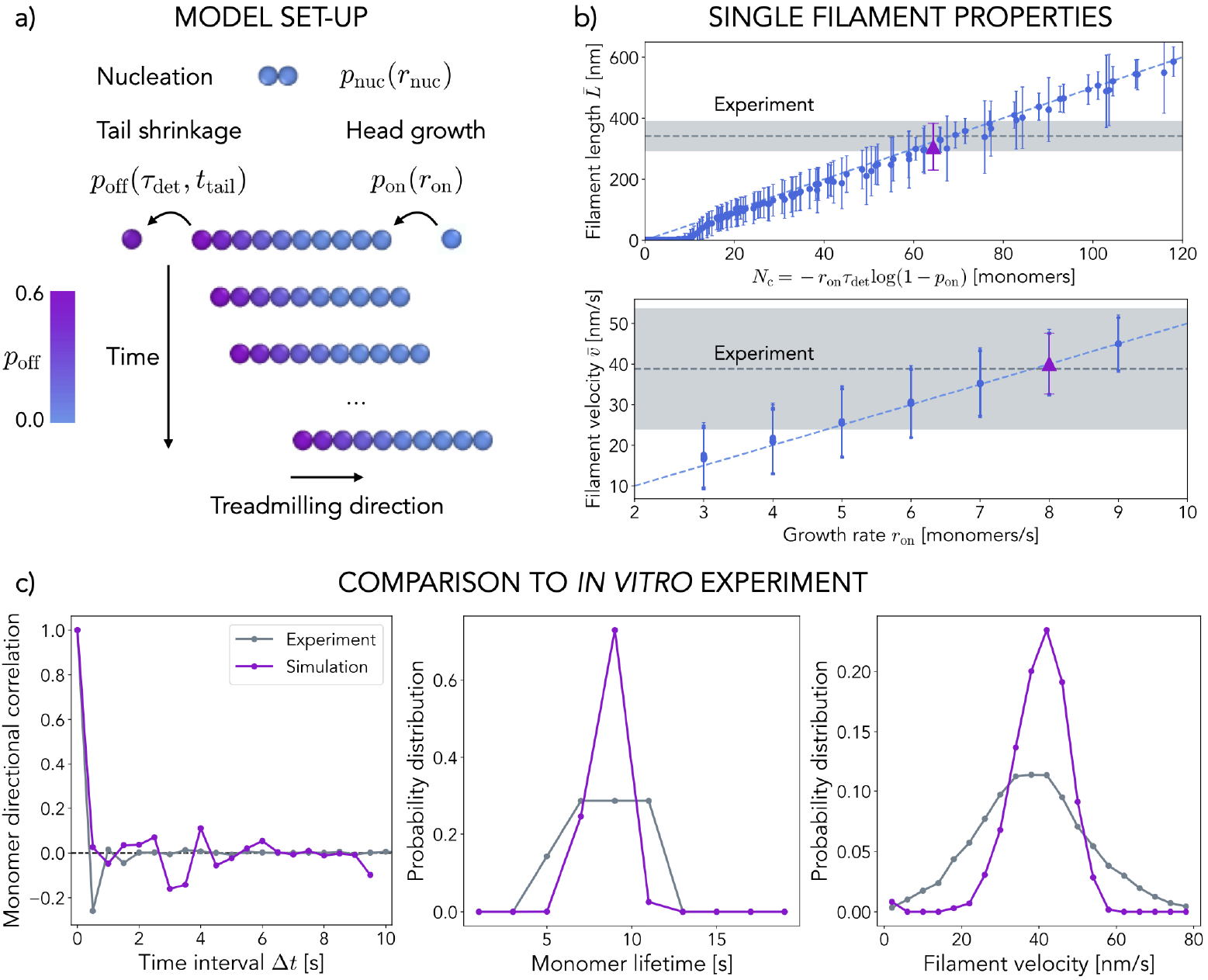
Coarse-grained model for treadmilling filaments. **a)** New filaments nucleate at a constant rate *r*_nuc_, grow with probability *p*_*on*_ = *r*_on_*dt*, and shrink with probability *p*_off_, determined by the monomer detachment time *τ*_det_ and the time the monomer has been in the system. This growth and shrinkage dynamics results in the apparent directional motion of the filaments over time. **b)** Top panel: Single filament length against the corresponding intrinsic size *N*_c_. The blue dashed line is the theoretical 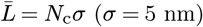. Bottom panel: Average treadmilling velocity 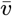 of a filament against the corresponding growth rate *r*_on_. The blue dashed line is the theoretical 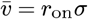. In both panels each point is the steady-state average over 20 replicas of single filament simulations with error bars for the standard error of the average. The grey dashed lines correspond to the fluorescence microscopy experiments of *in vitro* reconstituted *E. coli* FtsZ treadmilling filaments on supported lipid bilayers at 1.25*µ*M (shaded areas for error). Purple triangles correspond to *r*_on_ = 8 monomers/s and *τ*_det_ = 5 s, which fit experimental measurements best and are used for the simulation data in panel c. **c)** Single-filament dynamics in simulations (in purple) quantitatively matches *in vitro* FtsZ dynamics (in grey). Left panel: Directional autocorrelation of FtsZ monomers at increasing time intervals Δ*t* decays quickly, showing the static nature of individual monomers. Middle panel: Treadmilling filament velocity distribution. Right panel: Monomer lifetime distribution.

Treadmilling filaments are active, dissipating energy via nucleotide hydrolysis at the monomer-monomer interface. This results in a change in the interface domains of the monomers, which then tend to dissociate when they reach the tail of the filament [24, 28–30]. To capture these properties, we consider reactions at time intervals *dt*_react_ = 0.1 s during which individual filaments grow and shrink with probabilities *p*_on_ and *p*_off_ respectively, and new filaments are nucleated with probability *p*_nuc_. Tread-milling depends only on two parameters: the head polymerisation rate – *r*_on_ – and the typical time over which monomers become available for detachment as a result of hydrolysis – *τ*_det_. Here *r*_on_ (in monomers/s) captures both the bulk free monomer concentration, which we assume is constant, and the polymerisation rate constant, such that *p*_on_ = *r*_on_*dt*_react_. Monomers in solution are thus implicitly considered. Importantly, filament polymerisation is allowed only if free space for monomer addition is available around the filament head. Tail depolymerisation is modelled through 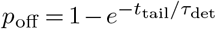, where *τ*_det_ (in s) effectively accounts for both the slow nucleotide hydrolysis at the monomer-monomer interface in the filament and the fast monomer dissociation at the filament tail, while *t*_tail_ is the time for which the tail monomer has been in the system. Figure S3 explores in depth the role of hydrolysis and dissociation reactions in depolymerisation kinetics. Given that monomers detach into solution upon depolymerisation, *p*_off_ is not dependent on the environment of the filament on the surface [31]. To reach steady state *p*_on_ = *p*_off_ must be satisfied, which defines an intrinsic size *N*_c_ = *−r*_on_*τ*_det_ log(1 *− p*_on_) around which filaments fluctuate while treadmilling at constant velocity *v*_*c*_ = *r*_on_*σ*. Note that this also defines a typical monomer life-time *T*_c_ = *−τ*_det_ log(1 *− p*_on_).

Finally, insertion of new filament nuclei into the system (modelled as dimers) is controlled by the imposed nucleation rate *r*_nuc_ (in nuclei/s), such that *p*_nuc_ = *r*_nuc_*dt*_react_. Like for the growth rate, *r*_nuc_ captures both the nucleation rate constant and the free monomer concentration.

## Single filament dynamics

We first perform simulations of individual filaments for a range of filament growth and detachment time parameter sets {*r*_on_, *τ*_det_ }. We initialise the system with a single filament nucleus and let it evolve in time (Supplementary Movie 1). As expected, filaments reach a steady-state where they fluctuate around a constant length and treadmilling velocity (Figure 1b), while the individual monomers display finite life-times (Figures S4-5). Experimentally, FtsZ protofilaments have been measured (*in vivo* and *in vitro*) to be between 100 and 500 nm long and to treadmill at speeds between 15 and 50 nm/s, while individual monomers turnover at lifetimes around 8 seconds, depending on the bacterial strain and conditions [25, 27, 32–34], all of which are well within the range we observe in simulations (Figure 1b).

We now turn to direct comparison of our model with experiments. For this purpose, we reconstituted *E. coli* FtsZ filaments (FtsA and fluorescently labeled FtsZ) on supported lipid bilayers (SLBs) in a reaction chamber with excess ATP/GTP and used total internal reflection fluorescence (TIRF) microscopy to image treadmilling dynamics and filament organization (see SI for further details on the experimental setup). Using previously developed image analysis software [35, 36] we measure filament lengths and velocities (grey dashed lines and shaded regions in Figure 1b, also reported in [35]) as well as individual monomer lifetimes (Figures S4-5). When we choose {*r*_on_, *τ*_det_ }model parameters that match this specific experimental system (the purple triangles in Figure 1b), the velocity and lifetime distributions also display an excellent agreement with the experimental data (Figure 1c, middle and right panels).

Importantly, the directional autocorrelation function of individual monomers for different time intervals Δ*t*, measured both in simulations and *in vitro* experiments, presents a sharp decay to zero (Figure 1c, left panel), indicating that individual monomers remain static even though the filament as a whole appears to be moving. The same is also clearly visible from the individual molecule track analysis *in vitro* (Figure S7). This lack of motility is an important feature of treadmilling filaments, which also suggests that modelling tread-milling as self-propulsion is likely unsuitable, as individual monomers do not display any directional motion. Consequently, pushing forces exerted by treadmilling filaments are negligible or small, as previously discussed for actin bundles [10]. Together, these results demonstrate that our model correctly captures the main aspects of treadmilling dynamics on biologically relevant time- and length-scales, that it can be benchmarked for a specific protein under particular conditions, and can be used for quantitative comparison with experiments.

## Collective filament dynamics

We are now in a unique position to explore the collective dynamics of many treadmilling filaments and how their mortality affects the formation of higher order structures. We nucleate filaments in the system, characterised by parameters {*r*_on_, *τ*_det_ }, at a constant nucleation rate of *r*_nuc_. The system generally reaches steady-state surface density *ρ* and average filament size 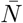 while the filament population undergoes continuous turnover (Figure S8). We observe three different collective regimes (Figure 2a), which depend exclusively on the treadmilling kinetics or the resulting intrinsic filament size *N*_c_ (Figures S8-10).

**FIG. 2.**
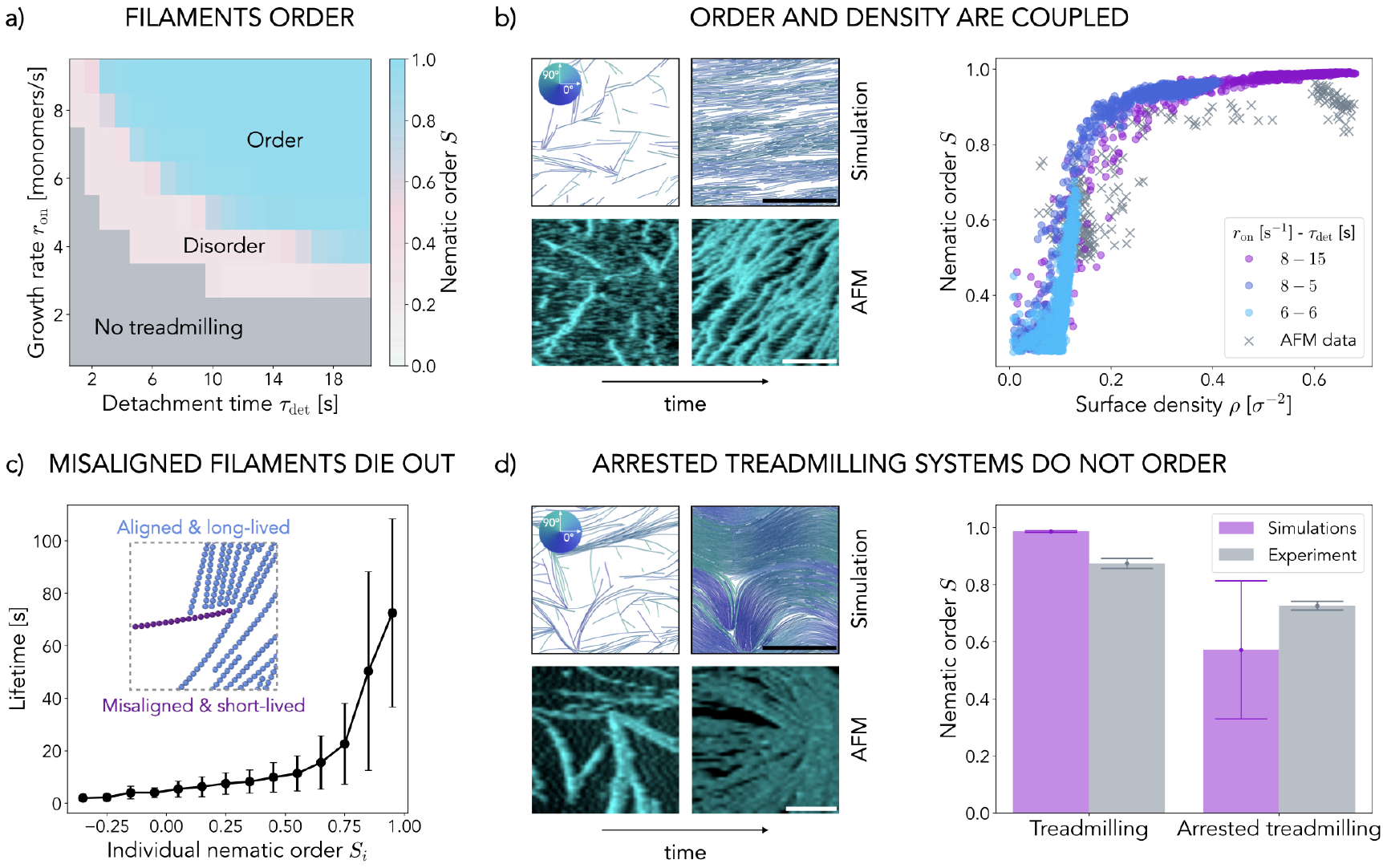
Collective behaviour of treadmilling filaments. **a)** Steady state nematic order parameter *S* measured in simulations (average over *N* = 10 replicas, *t >* 10 minutes). Grey indicates the quasi-empty systems of non-treadmilling filaments, blue the nematically-ordered system with polar bands, and pink the disordered transitional region. **b)** Left: Snapshots of treadmilling systems at early (left) and late (right) times in simulations (filaments coloured according to their orientation – see the colour wheel in the inset) and in HS-AFM experiment. Right: Nematic order *S* plotted against surface density *ρ* at different points in time for reconstituted FtZ on SLBs (AFM data — crosses) and simulations (coloured according to kinetics — circles). For experiments: data from *N* = 3 different movies is shown. For simulations: *N* = 10 different trajectories are shown for each parameter set. **c)** Filament lifetime for different levels of alignment (characterised by the individual nematic order *S*_*i*_). We consider only simulations for *N*_c_ *≥* 100 (*N* = 10 replicas per parameter set), which results in 226077 data points (filaments) in total binned in *S*_*i*_ with width 0.1. Inset: Snapshot of a *trapped* misaligned filament (purple) which dissolves over time (Movie S7). **d)** Left: Snapshots of systems with arrested treadmilling at early (left) and late (right) times in simulations (same kinetic parameters as panel b) and in HS-AFM experiments. Right: Steady state nematic order *S* for treadmilling and non-treadmilling systems. Bars and dots indicate the average for *t >* 600 s (*N* = 10 replicas in simulations) and error bars indicate the standard deviation. In experiment “treadmilling” corresponds to wild-type data and “arrested treadmilling” to the L169R mutant. Unless stated otherwise, simulation data corresponds to *r*_on_ = 8 s^*−*1^, *τ*_det_ = 15 s and *r*_nuc_ = 1 s^*−*1^ for *l*_p_ = 10 *µ*m and *D* = 100 nm^2^*/*s. Scale bars are 500 nm.

The first regime occurs for very low values of the intrinsic filament size *N*_c_ (grey area in Figure 2a), where the individual filaments die out rapidly (Figure S6), resulting in a quasi-zero surface density (see also Figures S8-10). As the intrinsic filament size increases (*N*_c_ *∼* 10) the system transitions into an unstable treadmilling regime, where filament size fluctuations are comparable to their size (Figure S6). The system is then populated by treadmilling filaments that stochastically emerge and disappear, never reaching high surface densities (the pink region in Figure 2a and Supplementary Movie 2). Finally, for higher values of *N*_c_ (*N*_c_ ≫ 10), the intrinsic size fluctuations become negligible compared to the filament size (Figure S6), and filaments enter a highly stable and persistent treadmilling regime in which they can achieve long lifetimes. This allows the system to build up its filament population and reach higher surface densities (Figures S8-10).

We find that the majority of this parameter space is characterised by the emergence of a large-scale nematic order (Figure 2a-b) and the formation of polar bands (snapshots in Figure 2a and Supplementary Movie 3). This behaviour is highly reminiscent of the *nematic laning* phase previously observed in self-propelled rods [37, 38] and is similarly characterised by high values of local and global nematic order parameter *S*, and high local but low global values of the polar order parameter *P* due to lanes formation (Figure S13). Unlike in self-propelled [8, 37, 38] or passive [39, 40] nematic systems, where the system density determines the ordering, in a treadmilling system the turnover kinetics (characterised by *N*_c_) dictates the emergent steady state ordering and density (Figures 2b and S10). We find that if monomer hydrolysis, and hence treadmilling, are arrested, and the filaments grow long, the system will evolve towards a highly populated but disordered configuration (Figures 2d and S15, see also Supplementary Movie 4). This occurs because, while thermal fluctuations and collisions can foster local alignment and bundling, the system cannot resolve the nematic defects that stochastically arise as filaments nucleate. Hence, treadmilling serves as an effective mechanism for defect healing.

We now turn to comparison of collective ordering with experiment. As shown in Figure 2b, High-Speed Atomic Force Microscopy (HS-AFM) images of *E. coli* FtsZ filaments on a supported bilayer bear a striking resemblance to simulation snapshots, with an evident transition to a high nematic order as the system becomes more populated (Supplementary Movie 5). These high resolution images allow us to quantify both the orientational order and the surface density of the system at any time from vector field analysis (Figure S16), again enabling a quantitative comparison with the simulations. Strikingly, Figure 2b demonstrates that both HS-AFM and simulation trajectories display the same coupling between orientational order and density, independently of the specific kinetic parameters used. Furthermore, when we perform HS-AFM with an FtsZ mutant with reduced GTPase activity and greatly inhibited depolymerisation (FtsZ L169R) [9], we observe that, just like in simulations, nematic defects are not resolved over time and the system remains at lower nematic order than wild type (Figure 2d and Supplementary Movie 6). These results indicate that the our modelling approach accurately describes FtsZ treadmilling behaviour as observed experimentally and should therefore provide insight into the mechanisms underlying the filament ordering.

In simulations we find that the ordering transition is driven by filaments growing against each other, which eventually selects for a common alignment direction, like in systems of self-propelled rods [37, 38, 41]. However, the underlying mechanism is completely different: unlike self-propelled rods that exert force when collided with another filament [42–45], treadmilling filaments stop growing when they reach another object, but still continue to depolymerise, which results in shrinkage, reorientation or even dissolution of the filaments. Ultimately, the fact that shrinkage is independent of growth causes misaligned filaments, which are more likely to grow against the neighbours than their aligned counterparts, to die out. Supplementary Movie 7 exemplifies a striking example of such a filament which, by being misaligned with its surroundings, stops growing and eventually dies.

Figure 2c measures the lifetime of each filament against its average alignment, characterised by the individual nematic order *S*_*i*_. These two variables are indeed highly correlated: only very aligned filaments (*S*_*i*_ *∼* 1) reach long lifetimes, up to several minutes, while strongly misaligned polymers die out after a few seconds on average and are gradually replaced with aligned ones. The aligned filaments also treadmill with a velocity that approaches their single-filament value *v*_c_ = *r*_on_*σ*, while the misaligned filaments typically have vanishing velocities (Figure S11). Globally this behaviour results in the progressive selection of the more stable filaments that align with the rest of the system, reminiscent of what was previously proposed for plant microtubules [46, 47]. The filament death and birth mechanism, and the establishment of order, are not rapid, but require turnover of a large number of filaments and typically take minutes (Figure S12).

## Treadmilling drives bacterial division ring condensation

To showcase the importance of the identified self-organisation mechanism in living cells, we investigate the role of treadmilling for bacterial division ring formation. Our *in vivo* experiments recently showed that treadmilling dynamics of FtsZ is essential for Z-ring formation in division of *B. subtilis* [23], but the details linking the two remain elusive. We now use our coarse-grained molecular model for treadmilling filaments to explore the underlying mechanism. Bacterial FtsZ has been observed to display mild natural curvature in live cells and *in vitro*, with diameters similar or smaller to typical cell diameters [13, 31, 48, 49]. In addition, its membrane anchor FtsA – which makes composite polymers with FtsZ – forms intrinsically curved filaments that therefore display a preference for curved membranes [50]. This suggests that the interplay between cell geometry and curvature of the (composite) filament should favour filament alignment along the circumference of the cell, similarly to what has been previously shown for MreB filaments [51–53]. We incorporate this curvature-sensing mechanism in the model, as shown in Figure 3a, by adding a circumference aligning force *f*_curv_ to the head and tail monomers of each filament (see SI for details). We find that a small force of this kind (*f*_curv_ = 5 *k*_B_*T/σ*) is sufficient to determine the global orientation of the filaments along the cell circumference (Figure 3a). These results point to the importance of the interplay between filament and cellular geometry in supra-molecular organisation.

**FIG. 3.**
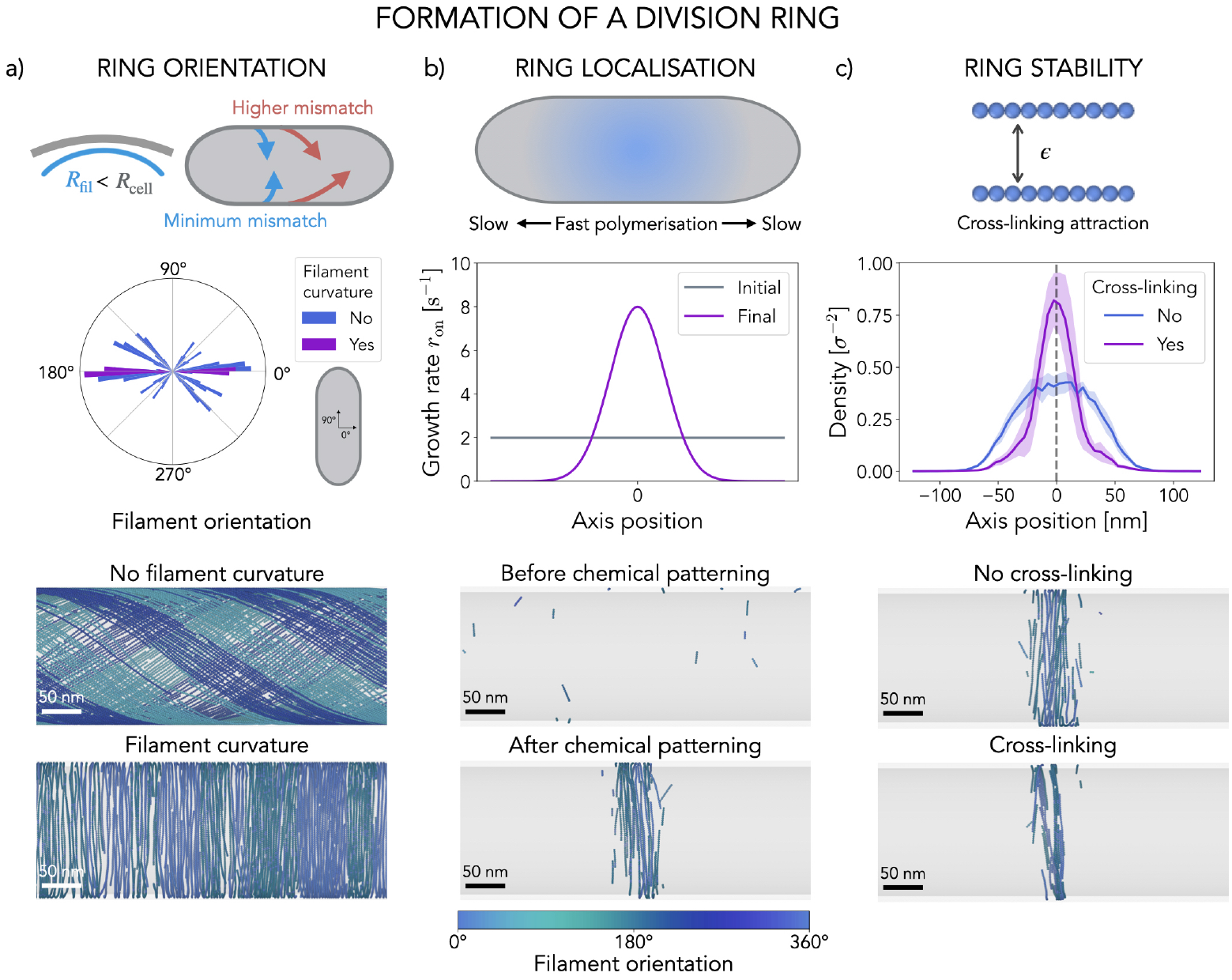
Formation of the bacterial division ring. **a)** Filament curvature and cell geometry drive the collective filament orientation along the cell circumference. Top panel: Illustration of the curvature-sensing mechanism of FtsZ/FtsA filaments. Middle panel: Distribution of filament orientations with and without filament curvature. Each bar plot corresponds to the average over *N* = 10 replicas in the nematic order region of the parameter space (*r*_on_ = 8 s^*−*1^, *τ*_det_ = 15 s). Bottom panel: Two representative snapshots of steady-state configurations of treadmilling filaments with and without filament curvature. **b)** Spatial modulation of FtsZ’s growth and nucleation kinetics mediates its midcell localisation. Top panel: Illustration of the typical FtsZ kinetics modulation observed *in vivo*. Middle panel: Example of a kinetics modulation combining an increase in the growth and nucleation rates around the midcell with a decrease at the poles. Bottom panel: Two representative snapshots of system configurations before and after the modulation of the kinetics is switched on. **c)** Attractive interactions stabilise the ring and mediate tight packing. Top panel: illustration of the cross-linking implementation. Middle panel: Late time (*t >* 8 minutes after the switch) density profiles along the cell axis with and without cross-linking interactions. Bottom panel: Two representative snapshots of steady-state rings with and without cross-linking interactions. In panels b and c we use switch parameters 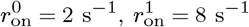 and *w*_prof_ = 100 nm for *τ*_det_ = 15 s. In all snapshots the filaments are coloured according to their orientation with respect to the cell circumference (see colour bar at the bottom) and the scale bars are 50 nm. For all simulations we set *f*_curv_ = 5 *k*_B_*T /σ, l*_p_ = 10 *µ*m and *D* = 100 nm^2^*/*s in a box of size *L* = 200*σ* = 1 *µ*m.

Bacteria have evolved a number of different molecular tools that act in complementary ways to position FtsZ filaments to the right division site – the midcell – at the right time. Such systems include dynamic chemical patterning [54–59], condensate formation [60] or nucleoid occlusion [61–64] and generally have the effect of spatially modulating FtsZ treadmilling kinetics, resulting in higher polymerisation rates at the midcell and lower ones around the poles (schematic in Figure 3b). Our model can incorporate this FtsZ growth modulation as an instantaneous change from a uniform distribution of growth and nucleation rates along the cell body to a Gaussian profile centered around the midcell for growth and nucleation rates, with a constant detachment time (Figure 3b, SI). We find that the resulting combination of the midcell enrichment and cell pole depletion of FtsZ proteins reliably localises the filaments to the midcell region (Figure 3b).

Finally, we consider one more model ingredient. It has been shown that FtsZ self-interaction, as well as cross-linking by FtsZ-binding proteins such as ZapA can promote Z-ring condensation, and that cross-linker deletion can lead to axial spreading of the rings [25, 35, 65, 66]. In line with this, we find that including explicit attractive interactions between filaments has an important stabilising effect on the ring structure, further promoting filament alignment and tight packing, and allowing for a sustained monomer accumulation in the ring over time (Figures 3c and S18). Taken together, these results show that treadmilling filaments in our model can spontaneously form stable ring-like structures when a combination of filament curvature, attractive inter-filament interactions, and spatio-temporal modulation of the filament kinetics are considered.

## Comparison with *in vivo* data

We are now in a good position to compare our model for Z-ring formation with live cell data. Our experiments have previously directly identified a very particular dynamics of Z-ring formation in living *B. subtilis* cells: localisation of the ring to the midcell, often referred to as ring condensation, occurs very rapidly, within 1 minute (Figure 4a), while ring maturation via growth and thickening occurs over several minutes (Figure 4b). If treadmilling is impaired early on, the Z-ring cannot form and remains diffuse, while impairing treadmilling during maturation disrupts proper division [23]. Our experiments also found that the average treadmilling velocity of filaments in mature Z-rings is substantially faster than in nascent ones (Figure 4c).

**FIG. 4.**
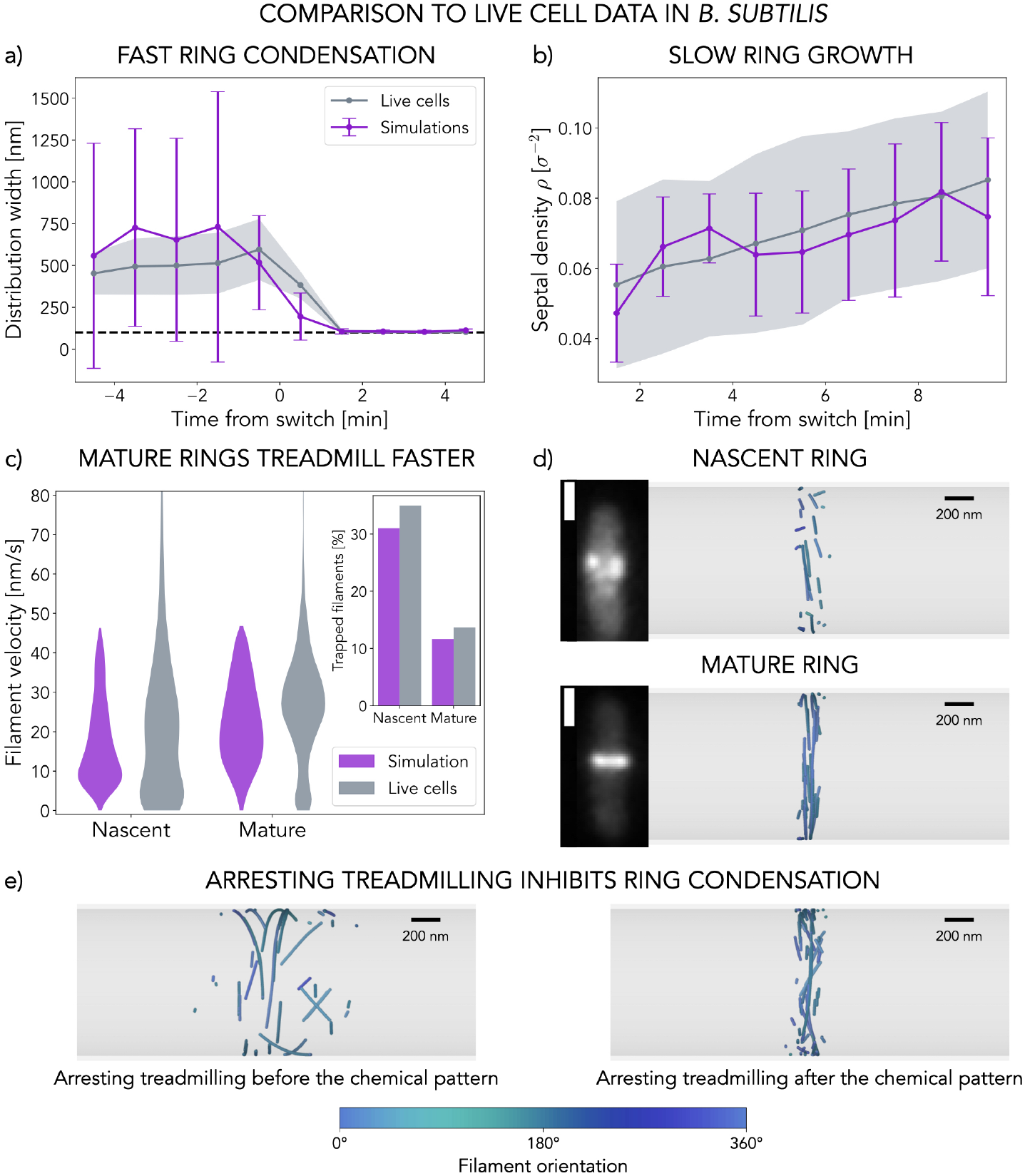
Z-ring formation *in vivo*. **a)** Z-ring condensation dynamics for live cells (grey) and simulations (purple). A rapid ring collapse around *t* = 0 minutes is observed. **b)** Ring density in time for live cells (grey) and simulations (purple). A positive average slope is observed, indicating the slow accumulation of proteins to the division site.**c)** Filament velocity distributions in nascent and mature rings for live cells (grey) and simulations (purple). Inset: Fraction of trapped filaments (*v <* 10 nm/s) for each distribution. In simulations we define *t <* 2 minutes for nascent and *t >* 8 minutes for mature rings. **d)** Representative snapshots of two successive ring configurations observed *in vivo* and in simulations. Microscopy images show fluorescently tagged FtsZ in live *B. subtilis* cells, scale bar is 1*µ*m. **e)** Representative snapshots of systems at *t* = 10 minutes after arresting treadmilling. Treadmilling is arrested 5 minutes before (left panel) and 1 minute after (right panel) the onset of the chemical pattern that modulates the growth kinetics. The simulation data in panels a), b) and c) corresponds to a switch with parameters 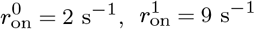 and 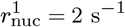 for *τ*_det_ = 15 s and *w*_prof_ = 100 nm. In panels a) and b) solid lines correspond to the average over samples (*N* = 10 replicas for simulations, *N* = 67 cells for experiments) and the shaded region or error bars to the standard deviation. Time *t* = 0 corresponds to the onset of the chemical pattern. Note that monomers in simulations are rendered with size 20 nm instead of the actual 5 nm for visualisation purposes and coloured according to their orientation with respect to the cell circumference (see the colour bar at the bottom). Scale bar is 200 nm.

To explore whether our model can capture the dynamics of Z-ring formation *in vivo*, and explain the mechanism behind it, we simulate systems on the cell scale (*L* = 3 *µ*m giving *R ∼* 1 *µ*m). We limit the total amount of monomers to the estimated amount of FtsZ in *B. subtilis* Z-rings *N*_max_ = 2000 (see SI for details), resulting in a relatively low monomer surface density. Strikingly, as shown in Figure 4a (see also Supplementary Movie 8), our model also displays rapid ring condensation, which occurs only a few seconds after the switch of the growth rate modulation profile. This behaviour is caused by the rapid dying out of the filaments in the growth-inhibited areas of the cell, as their depolymerisation rate is faster than the polymerisation rate. The whole process occurs on the scale of the single monomer’s lifetime, as controlled by *τ*_det_. Hence, treadmilling allows the system to rapidly respond to external chemical cues.

We further find, in agreement with what has been observed in live cell experiments, that after rapid ring condensation at the site of division, treadmilling filaments in the model accumulate rather slowly, over a time-span of several minutes. The simulations reveal a steadily increasing surface density of monomers in the midcell region that matches well the experimental measurements (Figure 4b). This feature is the inherent result of treadmilling-driven alignment: the ring structure transitions from an initially disordered state to an ordered state populated by several bundles of filaments tightly bound together (Figure 4d, Supplementary Movie 8) through the dissolution and replacement of misaligned filaments in a process that is analogous to what we also observed *in vitro* for *E. coli* FtsZ. Mature rings are thus composed of small patches of FtsZ organised in bands that are a few filaments wide and travel together, as previously observed in live-cell imaging [19, 67, 68] (Figure 4d and Supplementary Movie 8). These bands are tightly bound, displaying high local density of monomers, which is consistent with cryotomography experiments estimating *∼* 6 nm inter-filament distance in FtsZ rings [69].

Ring maturation is driven by the replacement of misaligned filaments, which is at heart a trial-and-error process, rendering this process slow. The decrease in the fraction of low-velocity, misaligned filaments in the mature ring also naturally leads to faster treadmilling velocities without individual filaments speeding up, which matches what has been measured in live cells (Figure 4c). In other words, the surviving filaments all run parallel to each other with treadmilling velocities close to maximal values. Finally, we found that arresting treadmilling in our model has similar effects to those observed *in vivo*. If treadmilling is arrested early on, before the onset of the chemical patterning, the filament population remains diffuse and never condenses to the midcell (Figure 4e, Supplementary Movie 9). If treadmilling is arrested after the switch of the chemical patterning, the ring forms at the midcell, but displays aberrant, “frozen”, configurations (Figures 4e and S21, Supplementary Movie 10). The crucial role of treadmilling in positioning of FtsZ ring that our model unveils has also been previously discussed in the context of *E. coli*: it’s been reported that mutants that are insensitive to classical positioning systems such as Min (FtsZ2, FtsZ9 or FtsZ100 for instance) all have very low or undetectable GTPase activity. This implicates that the lack of treadmilling underlies their inability to correctly position the ring [27].

Altogether, our treadmilling model naturally reproduces the key dynamical aspects of Z-ring formation observed in live cells. The FtsZ filaments are able to localise to the midcell on timescales similar to the monomer lifetimes, driven by the underlying spatial modulation of the polymerisation rates, while the tightly-packed division rings grow on substantially longer timescales by virtue of misaligned filaments dying and growing again until only the aligned ones remain. This mechanism is at play irrespective of the filament density: our simulations of treadmilling filaments on a planar surface constrained to *in vivo*-like low protein concentrations show that ordering still occurs and is driven by the same mechanism of death by misalignment (Figure S14). More generally, our model explains the importance of treadmilling – dynamic growth and shrinkage of filaments – in localising and timing bacterial division.

## Discussion

Here we developed a coarse-grained molecular model for explicitly treadmilling filaments in two dimensions, using a polymerisation kinetics that depends on two parameters only: the imposed growth rate *r*_on_ and the typical monomer detachment time *τ*_det_. Such polymers can grow and shrink on opposite ends at equal rates, virtually displaying directional motion while individual monomers remain static and undergo turnover. By tuning these two parameters the model can replicate experimentally observed FtsZ properties quantitatively, such as the filament lengths and treadmilling velocities, as well as the monomer lifetimes of treadmilling *E. coli* on supported bilayers. Since the model can capture a wide range of filament lengths and velocities by tuning the two parameters, it is also expected to be well-suited to study other cytoskeletal treadmilling proteins.

From a physics point of view, the explicit molecular nature of the model allowed us to flesh out a previously underappreciated or unrecognised mechanism behind self-organisation of treadmilling filaments – the nematic ordering via death of locally misaligned filaments. Our findings are experimentally confirmed by High-Speed AFM imaging of reconstituted FtsZ filaments which display the same ordering transition, while the individual monomers undergo turnover. The ordering transition, which shows quantitative matching between simulations and experiment, is completely controlled by the kinetics of this out-of-equilibrium system.

Our treadmilling model unveils a surprisingly rich collective behaviour arising from the dissolution and replacement of trapped filaments, which is characteristic of mortal polymers. These findings put forward treadmilling filaments as a paradigm for a dynamic, self-ordering system that is able to quickly remodel itself in response to external cues. Self-remodelling indeed allows treadmilling filaments to rapidly respond to external biases, such as local curvature or chemical patterning, which is not achievable in systems of static filaments. This suggests that both naturally occurring but also artificial treadmilling polymers could be a useful tool in areas such as synthetic cell development or programmable active matter. We showed that such dynamic structures are easily tunable, which should be experimentally testable, for example using microfluidic devices and chemical or photo patterning to generate different dynamic systems.

From a biological perspective, our study gives mechanistic insights into the molecular principles of rapid assembly of sparse cytoskeletal filaments into a dense Z-ring which, at least in *Bacillus subtilis* cells [23, 25], is required to recruit the septal cell wall synthesis machinery to mid-cell and thereby initiate bacterial cell division. We found that treadmilling-driven filament alignment, in tandem with filament sensing of cell curvature, cross-linking interactions, and spatiotemporal regulation of filament polymerization kinetics, drive Z-ring condensation – the rapid self-assembly of FtsZ filaments into a dense ring at the midcell. The onset of the process occurs on a timescale that is comparable to the single monomer lifetime while the further growth and maturation of the ring take minutes, as it requires stochastic dissolution and replacement of misaligned filaments. Quantitative matching between our simulations and live cell data confirms this picture.

Even in its simplicity, the model provides structural and dynamic insights into functional FtsZ assemblies and could be applied to unanswered questions in the field of bacterial division, such as the recruitment of bacterial wall synthesis machinery to the midcell, which ultimately divides the cell into two [15, 70, 71]. The model can help investigate how the dynamics of the Z-ring affects recruitment of the synthetic machinery and ultimately the growth of the cell wall at midcell.

Beyond their biological and biomimetic relevance, treadmilling filaments belong to a distinct class of active matter, which consumes energy not for motility but for turnover. This leads to a previously underappreciated mechanism of self-organisation and response to external biases, whereby activity allows to dissolve defects and collectively organise. This mechanism can also be expected to have further implications for filament collective behaviour, such as the response to perturbations and propagation of information, which will all be important to explore in the future.

## Acknowledgements

We thank Ivan Palaia and members of the Šarić lab for useful discussions, and Keesiang Lim and Richard W. Wong (WPI-Nano Life Science Institute, Kanazawa University) for providing access to HS-AFM. This work was supported by the Royal Society (Grant No. UF160266; C.V.C. and A.Š.), the European Union’s Horizon 2020 Research and Innovation Programme (Grant No. 802960; A.Š.), the Austrian Science Fund (FWF) StandAlone P34607 (M.L), a Wellcome Trust and Royal Society Sir Henry Dale Fellowship (Grant No. 206670/Z/17/Z; S.H. and K.D.W.). For the purpose of open access, the authors have applied a CC BY public copyright licence to any Author Accepted Manuscript version arising from this submission.

## Code and data availability

Appropriate documentation and example files to replicate the simulation results presented in this work together with all necessary analysis code can be found at the following [public repository](https://doi.org/10.5522/04/24754527) [72]. Additionally, a maintained version of the code is available on [GitHub](https://github.com/cvanhille/treadmilling_filaments) [73].

