## Supplementary Information for "Self-organisation of mortal filaments: the role of FtsZ treadmilling in bacterial division ring formation"

[1] *Institute of Science and Technology Austria, Klosterneuburg, Austria**
[2] *Department of Physics and Astronomy, Institute for the Physics of Living Systems, University College London, London, United Kingdom*
[3] *Centre for Bacterial Cell Biology, Biosciences Institute, Newcastle University, Newcastle upon Tyne, United Kingdom*
[4] *School of Life Sciences, The University of Warwick, Coventry, United Kingdom*
(Dated: March 13, 2024)

This Supplementary Information provides additional details on the treadmilling model presented in this work together with technical information about its implementation in simulations. It also includes complementary supporting data for the results presented in the manuscript as well as details on some of the measurements and experimental setup.

#### A. Simulation details

We consider coarse-grained polymers made of spherical beads with diameter $\sigma = 5$ nm in a two-dimensional box of size $L$. Filament stretching rigidity is captured by harmonic springs between pairs of neighbouring beads: $E_{\mathrm{bond}}(r) = K_{\mathrm{bond}}(r-\sigma)^2$. Filament bending rigidity is captured by harmonic angle potentials between monomer triads: $E_{\mathrm{bend}}(\alpha) = 0.5K_{\mathrm{bend}}(\theta-\theta_0)^2$ where $\theta$ is the angle formed by the three monomers involved and $\theta_0 = \pi$ is the straight equilibrium configuration. Note that we can then define the persistence length of the filaments as $l_{\mathrm{p}} = 2K_{\mathrm{bend}}\sigma/k_{\mathrm{B}}T$. Typically we set $K_{\mathrm{bond}} = 1000\ k_{\mathrm{B}}T/\sigma$ and $K_{\mathrm{bend}} = 1000-10000\ k_{\mathrm{B}}T$ (so filaments are quite stiff — $l_{\mathrm{p}} > 1\ \mu$m). Additionally, to model the cross-linking interactions mediated by partner proteins such as ZapA, we can introduce a Lennard-Jones potential between monomers of different filaments. This potential is of the form $E_{\mathrm{LJ}}(r) = 4\epsilon[1/r^{12} - 1/r^6 - (1/r_{\mathrm{c}}^{12} - 1/r_{\mathrm{c}}^6)]\ \forall\ r < r_{\mathrm{c}}$ and $E_{\mathrm{LJ}}(r) = 0\ \forall\ r > r_{\mathrm{c}}$, which is shifted to guarantee continuity at $r = r_{\mathrm{c}}$. Note that, because we expect cross-linking interactions to be strong and short-ranged we choose a cutoff at distance $r_{\mathrm{c}} = 1.5\sigma$ and a large interaction strength $\epsilon = 24\ k_{\mathrm{B}}T$. When no cross-linking effects are present we implement volume exclusion via hard sphere interactions between beads. Finally, because FtsZ filaments are curved and live on a curved surface a tension will arise from the mismatch between the filaments intrinsic curvature and the one they sense and have to adapt to on the surface. Note that this sensed curvature will depend on their orientation, such that if they align with the cell axis filaments will sense a flat surface but if they align with the circumference they will sense a curvature $c_{\mathrm{surf}} = 1/R_{\mathrm{cell}}$. FtsZ filaments (and especially FtsZ/FtsA composite filaments) are typically more curved than the cell (or at least similarly curved) [1–5] such that this curvature tension should result in an effective force that aligns filaments along the circumference of the cell. When simulating cell-like systems, therefore, we introduce an additional force $f_{\mathrm{curv}}$ on the head and tail monomers of each filament to align it with the circumference direction. The $X$ and $Y$ components of this force take the form $f_X = f_{\mathrm{curv}}\ (1.0-\cos(\alpha))\ \sin^2(\alpha)\cos(\alpha)$ and $f_Y = -f_{\mathrm{curv}}\ (1.0-\cos(\alpha))\ \sin(\alpha)\cos^2(\alpha)$, where $\alpha$ is the angle of the tail-to-head vector with the circumference axis ($X$ axis of the simulation box). Periodic boundary conditions (PBCs) are implemented in all directions.

We evolve our model in Molecular Dynamics (MD) to capture the correct diffusive dynamics of proteins at this scale. We integrate the Langevin equation of motion in time over time steps of size $dt_{\mathrm{MD}} = 0.001\tau$ ($\tau$ being the simulation time unit) at constant temperature $T = 1$ (in reduced units). On top of this we implement growth, nucleation and shrinkage reactions according to our treadmilling kinetics model over reaction steps of size $dt_{\mathrm{react}} = 0.1$ s. The following reactions are imple-

*

mented in the system: 1) Nucleation: New filaments are nucleated in the system at a rate $r_{\rm nuc}$ s$^{-1}$. A new filament dimer is created at a random position and orientation with probability $p_{\rm nuc} = r_{\rm nuc} dt_{\rm react}$ as long as no overlaps forbid it. 2) Growth: New monomers are added to the head of existing filaments at a rate $r_{\rm on}$ s$^{-1}$. A new monomer is added to the system, at the equilibrium position given by the current position and orientation of the existing filament, with probability $p_{\rm on} = r_{\rm on} dt_{\rm react}$ as long as no overlaps forbid it. 3) Shrinkage: Tail monomers are removed from the system with probability $p_{\rm off}(t_{\rm tail}, \tau_{\rm det}) = 1 - e^{-t_{\rm tail}/\tau_{\rm det}}$, where $t_{\rm tail}$ is the time since it was added to the filament, irrespective of the surrounding conditions. We consider that, unlike for growth, this shrinking probability should be independent of the local conditions (i.e. crowding) because monomers detach into solution upon depolymerisation and therefore need no space on the plane to diffuse away [2]. Finally, if at depolymerisation the filament consists only of two monomers then the whole dimer is removed from the system. Note that, because we are mixing molecular dynamics and polymerisation kinetics two different timescales emerge in our system. On one hand, the characteristic time of the kinetics is set by our reaction time step $dt_{\rm react} = 0.1$ s, which allows us to span physiologically relevant growth rates $r_{\rm on}$ while optimising reaction frequency. On the other hand, the diffusive dynamics of the particles allow us to define a mapping between simulation time $\tau$ and real time $t$ that sets the value of the diffusion coefficient of monomers $D$ (in nm$^2$/s) to physiologically relevant values (see Figure S1). All the simulations are run using a custom version of the LAMMPS (Large-scale Atomic Molecular Massively Parallel Simulator) Molecular Dynamics package [6] available on GitHub [7]. Reactions, in particular, which involve the creation and deletion of particles in the system are implemented through a modified version of the bond/react fix in LAMMPS [8, 9]. Appropriate documentation and example files to replicate the results presented in this work can be found on the following public repository [10]. Additionally, a maintained version of the code is available on GitHub [7].

#### B. Treadmilling polymerisation kinetics and their molecular origin

Treadmilling is a type of polymerisation kinetics whereby filaments grow and shrink on opposite ends at a constant rate, by addition and removal of monomers. This is an active process involving energy dissipation via GTP hydrolysis and a structural transition of the monomer upon polymerisation from a relaxed or solution conformation to a tense or filamentous form (also known as the cytomotive switch) [11, 12]. During nucleotide hydrolysis FtsZ monomers change their interface properties and hence their dissociation constant $K_{\rm d}$, such that GTP-bound molecules tend to form filaments while GDP-bound molecules tend to dissociate back to the cytosol (see Figure S2a) [13]. Additionally, FtsZ monomers in solution or relaxed form are asymmetric, such that interface formation is easier on one end of the filament than the other, which results in faster exchange kinetics at the head of the filament than at the tail, as depicted in Figure S2b and already proposed by Wegner in 1976 [14]. The combination of asymmetric exchange rates with nucleotide hydrolysis then enables the net growth of filaments from the head by association of GTP-bound monomers and their shrinking from the tail by dissociation of GDP-bound monomers that accumulate towards the tail as nucleotide hydrolysis takes place at the monomer-monomer interface (see Figure S2c) [11, 12, 15, 16]. Note that because tense GTP-bound monomers in the core of the filament are stabilised on both sides by their neighbours while the tail monomer is only stabilised on one interface, fragmentation is expected to be quite rare and dissociation mainly occurs at the tail. In summary, the combination of asymmetric polymerisation kinetics and GTP hydrolysis in FtsZ result in treadmilling filaments that grow and shrink on opposite ends at rates that are controlled by the complex balance between association, dissociation and nucleotide hydrolysis and exchange reactions. A very comprehensive model of this dynamics has been proposed by Corbin and Erickson [16]. While such a detailed model is too complex to consider in our case, where we also need to explicitly model multiple filaments that interact with one-another, we can draw inspiration to design a simplified version of treadmilling kinetics. In the end, the main elements of treadmilling are 1) a net filament growth rate, which depends on the monomer concentration in solution, the amount of GTP and the relaxed to tense transition; 2) a typical timescale after which monomers in the filament tend to dissociate from the tail, which depends on the nucleotide hydrolysis rate and the dissociation reaction of GDP-bound monomers; and 3) a filament nucleation rate, which again depends on monomer concentration, GTP and relaxed to tense transitions, but need not be the same as the growth

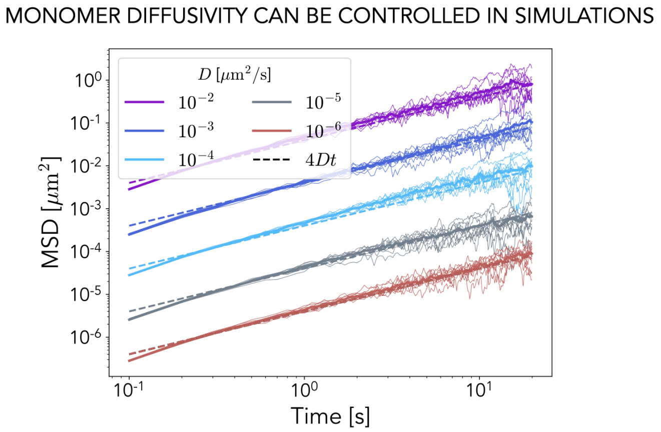


FIG. S1. **Controlling monomer diffusivity in simulations** Mean square displacement (MSD) of individual monomers over time for different simulation parameters that each result in a different and precisely controlled diffusion coefficient $D$, as illustrated by the dashed lines. $N = 10$ replicas for each value of $D$.

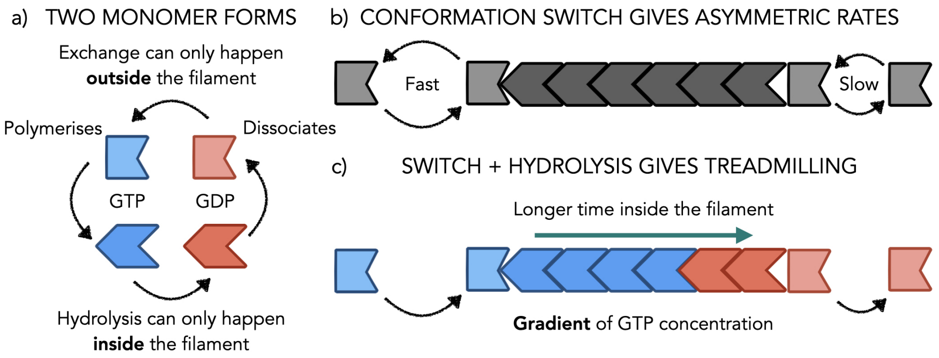


FIG. S2. **Treadmilling kinetics a)** Nucleotide state and monomer conformation schematics **b)** Polymerisation rates asymmetry arising from the conformation switch of monomers upon polymerisation. **c)** Treadmilling with asymmetric polymerisation and hydrolysis of nucleotides resulting in growth and shrinkage on opposite sides with the emergence of a GTP to GDP concentration gradient from head to tail.

rate. These are therefore the only three parameters we will keep in our simplified model for treadmilling. Note for instance that the fragmentation rate estimated by Corbin and Erickson is several orders of magnitude lower than any other rate in the system, which is why we also choose to neglect this effect in our approach.

### C. A simplified model for treadmilling kinetics

Let us further discuss our simplified model of treadmilling kinetics. In our approach we control the growth-shrinking kinetics of filaments via only two free parameters: the imposed growth rate $r_{\text{on}}$ and the detachment timer $\tau_{\text{det}}$. The first ($r_{\text{on}}$) corresponds to effective rate at which filaments grow, which englobes both the reaction rate constant and concentration effects. Note that we generally assume well-mixed conditions and do not consider the solution concentration of FtsZ independently. The second parameter ($\tau_{\text{det}}$), on the other hand, controls the time after which monomers in the filament become available for dissociation (removal).

In reality, the depolymerisation of tail monomers involves two reactions: hydrolysis of the associated nucleotide (GTP to GDP) and dissociation of the monomer after hydrolysis, each controlled by rates $r_{\text{hyd}} = 1/\tau_{\text{hyd}}$ and $r_{\text{dis}}$ respectively (Figure S3a). The overall shrinking rate $r_{\text{off}}$ will therefore result from a combination of the two. Let us now explore in detail how $r_{\text{off}}$ should depend on $\tau_{\text{hyd}}$ and $r_{\text{dis}}$ and

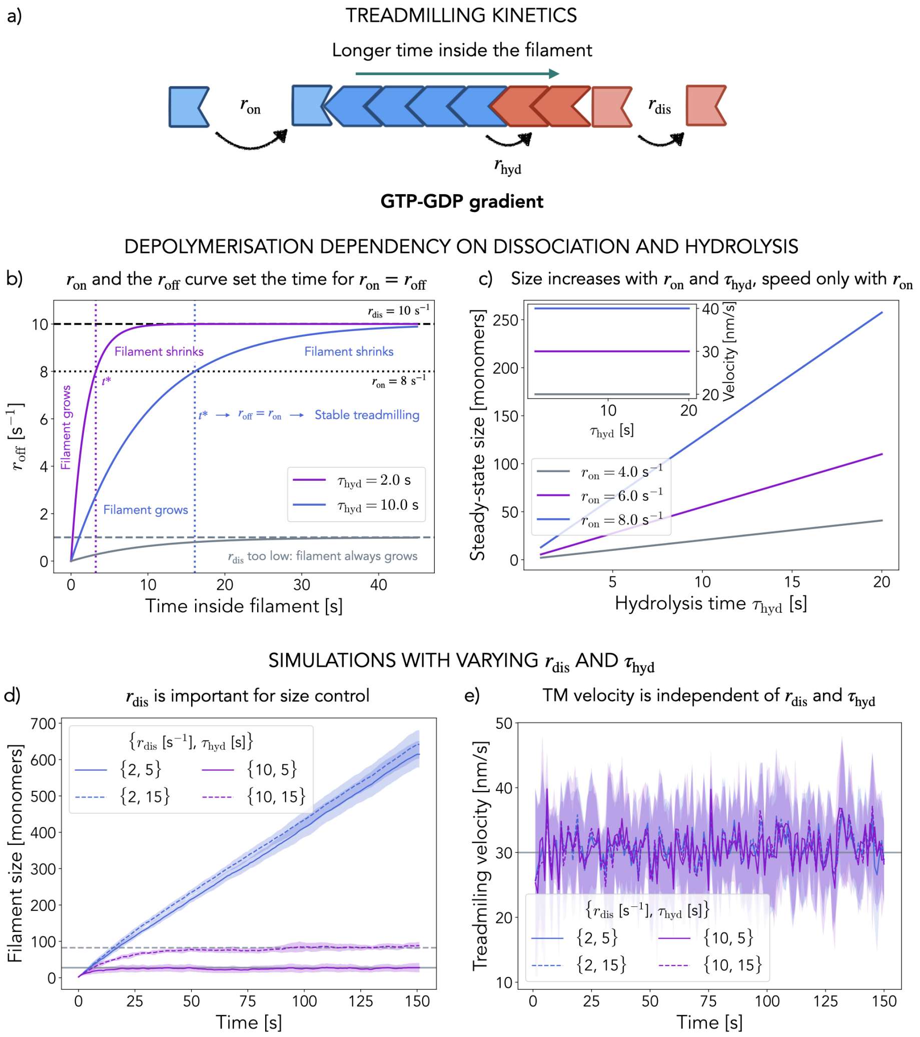


FIG. S3. **Treadmilling kinetics a)** Schematic of how the hydrolysis and dissociation reactions are involved in filament tail depolymerisation. **b)** $r_{off}$ is not constant and varies with the time a monomer has spent in the filament. This variation is controlled by $r_{dis}$ and $\tau_{hyd}$: $r_{off}$ relaxes to $r_{dis}$ over $\tau_{hyd}$. If $r_{on} = 8$ s$^{-1}$ (dotted black line) then only at $t^*$ (vertical dotted lines) are the two rates equal. The value of $t^*$ depends on $\tau_{hyd}$. **c)** Steady-state filament size and treadmilling velocity for different $\tau_{hyd}$ and $r_{on}$. Only the filament size depends on both while the velocity depends only on the growth rate. **d-e)** Filament size (d) and treadmilling velocity (e) over time for single treadmilling filaments in our Molecular Dynamics model, varying $r_{dis}$ and $\tau_{hyd}$ for a fixed $r_{on} = 6$ s$^{-1}$. Only filaments with large $r_{dis}$ show stable treadmilling at a constant size. This size depends on $\tau_{hyd}$. Treadmilling velocity is stable for all parameters and only depends on $r_{on}$. Gray lines in panel d) correspond to the expected steady-state size for $\tau_{hyd} = 5$ s (solid) and $\tau_{hyd} = 15$ s (dashed), assuming $r_{dis} > r_{on}$. Lines are the average over $N = 10$ replicas for each parameter set and the shaded regions correspond to the standard deviation.

justify how we simplify our model to control it via a single free parameter $\tau_{det}$. The probability for a tail monomer to detach from the shrinking end over a time interval $dt$ will be $p_{off} = r_{off}\ dt = p_{hyd}\ p_{dis}$, where $p_{hyd}$ is the probability that the tail monomer has hydrolysed its nucleotide and $p_{dis}$ the probability that such a monomer dissociates over the time interval. We can then write $p_{dis} = r_{dis}\ dt$ with $r_{dis}$ the dissociation rate. Hydrolysis of the nucleotide can only happen within the filament, as it required the full monomer-monomer interface and, because all non-hydrolysed interfaces in the polymer are the same, should occur at a constant hydrolysis rate $r_{\rm hyd} = 1/\tau_{\rm hyd}$. As a result, the probability that a monomer's nucleotide has been hydrolysed will increase over time as $p_{\rm hyd}(t) = 1 - \exp(-t/\tau_{\rm hyd})$, where $t$ is the time the monomer has spent in the filament. Putting everything together, we get that the depolymerisation rate $r_{\rm off}$ generally depends on $r_{\rm dis}$ and $\tau_{\rm hyd}$ as $r_{\rm off}(t) = r_{\rm dis}\,[1 - \exp(t/\tau_{\rm hyd})]$.

In summary, hydrolysis renders the depolymerisation rate time-dependent, increasing from 0 to $r_{\rm dis}$ over a typical time $\tau_{\rm hyd}$ as the time monomers spend in the filament increases. Consequently, as we show in Figure S3b given a growth rate $r_{\rm on}$, there will only be one time $t*$ for which the two rates become equal ($r_{\rm off}(t*) = r_{\rm on}$) and stable treadmilling is achieved. For shorter times $r_{\rm off}(t < t*) < r_{\rm on}$ and the filament grows and for longer times $r_{\rm off}(t > t*) > r_{\rm on}$ and the filament shrinks. Note that this time dependency of the depolymerisation rate, which results from the hydrolysis, is a hallmark of treadmilling, allowing for filament length control [17]. Importantly, a sufficiently large value of $r_{\rm dis}$ is required for stable treadmilling, as $r_{\rm dis} < r_{\rm on}$ implies $r_{\rm off}(t) < r_{\rm on}$ for all times. Assuming dissociation is sufficiently fast then, we expect the filament size will be controlled by both $r_{\rm on}$ and $\tau_{\rm hyd}$ while the treadmilling speed will be determined solely by the growth rate $r_{\rm on}$ (Figure S3c).

This analysis is consistent with the current literature on FtsZ, where the hydrolysis has been estimated to be much slower than the dissociation ($\tau_{\rm hyd} \gg 1/r_{\rm dis}$). Corbin and Erickson fit these parameters to $\tau_{\rm hyd} = 2.5 - \infty$ s and $r_{\rm dis} = 6.5$ s$^{-1}$ respectively [16] while Loose and Mitchison measure $\tau_{\rm hyd} = 12$ s and $r_{\rm dis} = 13.25$ s$^{-1}$ [1]. According to this picture then, given that hydrolysis is slow, it would take a long time for FtsZ monomers to hydrolyse their nucleotide and become available for detachment, but the depolymerisation rate would still be high, matching the growth rate $r_{\rm on}$.

Taking all this into consideration and striving for simplicity in our model, we decided to retain only one parameter to control depolymerisation: $\tau_{\rm det}$, the typical time over which monomers become available for detachment. We implement it into our model through the probability for removing a tail monomer over timesteps of size $dt = 0.1$ s: $p_{\rm off}(t_{\rm tail}, \tau_{\rm det}) = 1 - e^{-t_{\rm tail}/\tau_{\rm det}}$, where $t_{\rm tail}$ is the time since it was added to the filament. In this way, the depolymerisation rate retains the time-dependency resulting from hydrolysis, as discussed above. Note that $\tau_{\rm det}$ corresponds to $\tau_{\rm hyd}$ for fast dissociation ($r_{\rm dis} = 10$ s$^{-1}$) and can be related to $\tau_{\rm hyd}$ and $r_{\rm dis}$ by equating the time $t^*$ for stable treadmilling in both cases, yielding: $\tau_{\rm det}\log(1 - r_{\rm on}/10) = \tau_{\rm hyd}\log(1 - r_{\rm on}/r_{\rm dis})$.

Nonetheless, for the sake of completeness, in Figure S3d-e we take a step further and implement the two depolymerisation parameters – $r_{\rm dis}$ and $\tau_{\rm hyd}$ – explicitly in our Molecular Dynamics model, adding another level of complexity. The simulations are carried out in the same conditions as described previously for single filament simulations (Figure 1 of the main manuscript), only now $p_{\rm off}(t_{\rm tail}, r_{\rm dis}, \tau_{\rm hyd}) = r_{\rm dis}\,(1 - e^{-t_{\rm tail}/\tau_{\rm hyd}})\,dt$. All the data shown in Figure S3d-e corresponds to a fixed growth rate $r_{\rm on} = 6$ s$^{-1}$. Consistent with our predictions from above, only when the dissociation rate $r_{\rm dis}$ is large do these filaments achieve stable treadmilling where the filament size fluctuates around a constant value which in turn depends on $\tau_{\rm hyd}$. Otherwise, the depolymerisation rate is always too low and the filaments grow indefinitely. Note as well that the treadmilling velocity – measured as the filament head velocity – is independent of both $r_{\rm dis}$ and $\tau_{\rm hyd}$, as expected. Altogether, these results and analysis showcase that our model simplification, where the filament depolymerisation only depends on the time it takes for filaments to become available for detachment ($\tau_{\rm det}$), is a sensible one.

#### D. Single filament analysis

To test the validity of our modelling approach we simulate single filaments treadmilling in a box of size $L = 400\sigma$ for a wide range of parameters $\{r_{\rm on}, \tau_{\rm det}\}$ over $t_{\rm max} = 300$ seconds to guarantee proper steady-state sampling of their dynamic properties. The structural parameters of the filaments are kept constant at $K_{\rm bond} = 10^3 k_{\rm B}T$ and $K_{\rm bend} = 10^5 k_{\rm B}T$, thus simulating stiff and straight filaments. Simulations are initiated with a single filament nucleus in the center of the box and no new dimer nucleation is considered ($r_{\rm nuc} = 0$ s$^{-1}$). We characterise the steady-state filament size and its fluctuations over the last 2 minutes of simulation ($t \in (180, 300)$ seconds): $\bar{N} = \langle N_t \rangle_{t\in(180,300)}$, $\sigma_N^2 = \langle (N_t - \bar{N})^2 \rangle_{t\in(180,300)}$. In Figure S4a-b we present how the intrinsic size $N_{\rm c} = -r_{\rm on}\tau_{\rm det}\log(1 - p_{\rm on})$ and the steady-state size $\bar{L} = \sigma\bar{N}$ each depend on the model parameters. Note that they display a similar dependency and, as shown in Figure 1 of the main text the two variables collapse onto each other, showing that the steady-state size of treadmilling filaments can be precisely controlled by the two parameters $r_{\rm on}$ and $\tau_{\rm det}$. In Figure S6 we illustrate how the rescaled

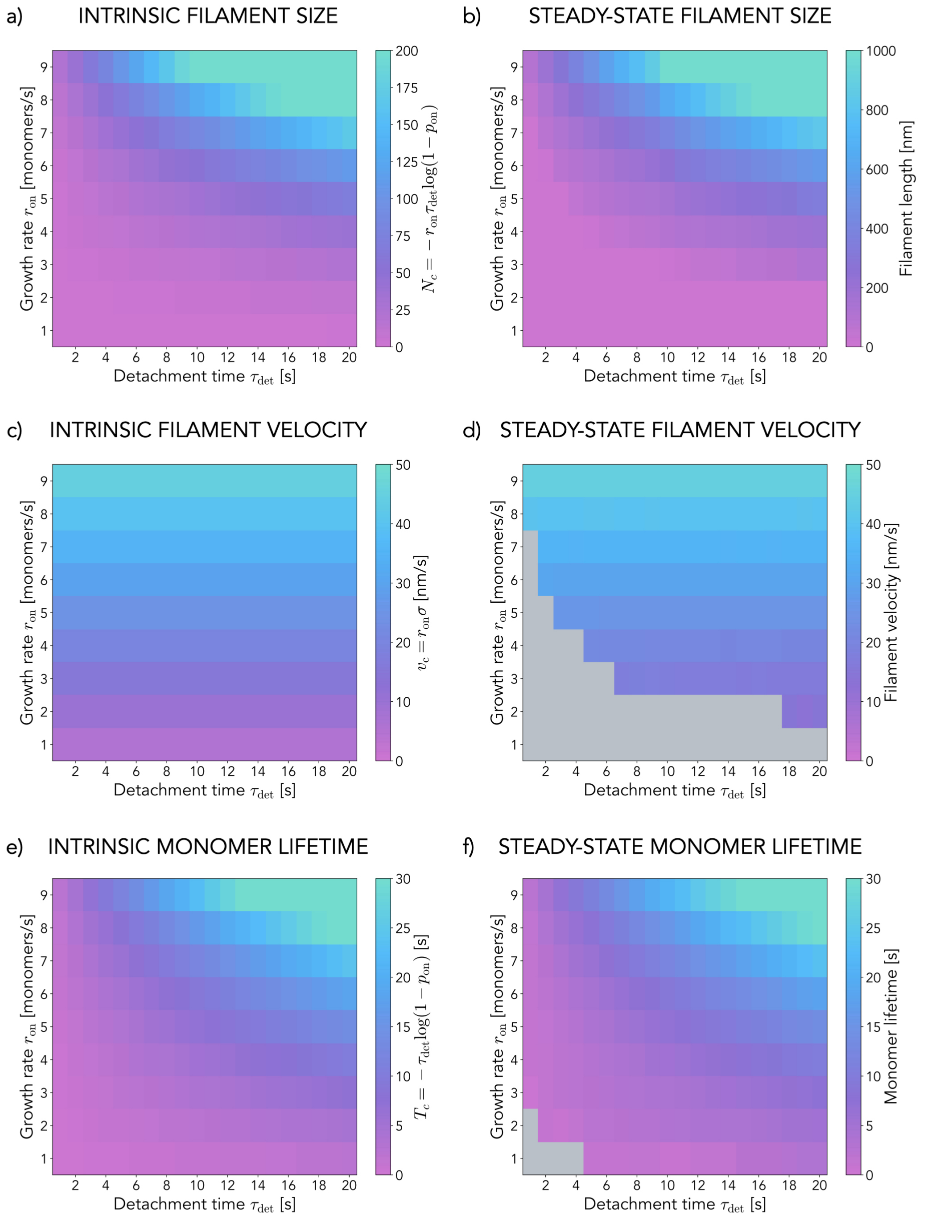


FIG. S4. **Single filament properties a)** Intrinsic filament size $N_{\rm c}$ dependency on the two model parameters $\{r_{\rm on}, \tau_{\rm det}\}$. **b)** Steady-state filament length dependency on the two model parameters $\{r_{\rm on}, \tau_{\rm det}\}$. **c)** Intrinsic filament velocity $v_{\rm c}$ dependency on the two model parameters $\{r_{\rm on}, \tau_{\rm det}\}$. **d)** Steady-state filament velocity dependency on the two model parameters $\{r_{\rm on}, \tau_{\rm det}\}$. **e)** Intrinsic monomer lifetime $T_{\rm c}$ dependency on the two model parameters $\{r_{\rm on}, \tau_{\rm det}\}$. **f)** Steady-state monomer lifetime dependency on the two model parameters $\{r_{\rm on}, \tau_{\rm det}\}$. Values in panels **b)**, **d)** and **f)** correspond to the average over 20 replicas for each parameter set.

size fluctuations $\sigma_N/\bar{N}$ are controlled by the intrinsic size $N_{\rm c}$, such that as $N_{\rm c}$ increases treadmilling becomes more stable and fluctuations are negligible compared to the filament size. Similarly, we also characterise the steady-state velocity of filaments as the average displacement of the filament head (again over the last two minutes of simulation): $\bar{v} = \langle dr_{\rm head}/dt \rangle_{t\in(180,300)}$ which, as shown in Figure S4c-d and Figure 1 of the main text only depends on the imposed growth rate as $\bar{v} = v_{\rm c} = r_{\rm on}\sigma$. Finally, we can also measure the average monomer lifetime over the course of the simulation $\bar{T} = \langle t_{\rm depol} - t_{\rm pol} \rangle$ which again can be predicted from the model parameters as $T_{\rm c} = -\tau_{\rm det} \log(1 - p_{\rm on})$, as illustrated by Figure S4e-f and in Figure S5. All statistics are performed over $N = 20$ different replicas of the system for each parameter set $\{r_{\rm on}, \tau_{\rm det}\}$.

#### E. Collective dynamics analysis

We characterise the collective behaviour of treadmilling filaments in systems of size $L = 200\sigma$ in which polymers with $K_{\rm bond} = 10^3 k_{\rm B}T$ and $K_{\rm bend} = 10^3 - 10^4 k_{\rm B}T$ ($l_{\rm p} = 10 - 100\mu$m) are nucleated in the form of dimers at constant rates $r_{\rm nuc} = 1$ s$^{-1}$ or $r_{\rm nuc} = 5$ s$^{-1}$ and evolve under kinetic parameters $\{r_{\rm on}, \tau_{\rm det}\}$ for $t_{\rm max} = 20$ minutes. Because we are comparing with High Speed Atomic Force Microscopy data from reconstituted *E. coli* FtsZ on supported lipid bilayers (SLBs), for which the monomer diffusion coefficient was estimated to be around $D = 10^{-4}$ $\mu$m$^2$/s [18], in our simulations we also set $D = 100$ nm$^2$/s. As exemplified in Figure S8, all relevant system variables stabilise around constant values after a few minutes, indicating that treadmilling systems generally relax to a well-defined collective steady state characterised by constant values of the surface density, filament number and average size and nematic and polar order parameters $S$ and $P$ respectively. We characterise this steady-state for different kinetic parameters by averaging the relevant variables in time over the last 10 minutes of simulation. We define the surface density $\rho_t = M_t/L^2$ where $M_t$ is the total number of monomers in the system at time $t$. Similarly, we define $S_t = 0.5(3\langle(\mathbf{u}_i\mathbf{u}_j)^2\rangle_{\{i,j\}} - 1)$ and $P_t = \langle \mathbf{u}_i\mathbf{u}_j \rangle_{\{i,j\}}$, where $\mathbf{u}_i$ is the unit vector along the direction of bond $i$, $\langle\rangle_{\{i,j\}}$ indicates the average over all pairs of bonds and the subscript $t$ indicates the time dependence of $S$ and $P$.

As indicated in Figure 2 of the main text, we find that treadmilling systems in the right kinetic regime will spontaneously organise into dynamic structures with high nematic order and surface density. This transition depends only on the kinetic parameters $r_{\rm on}$ and $\tau_{\rm det}$, and appears to be independent of the nucleation rate $r_{\rm nuc}$, as shown in Figure S9. Indeed, we find that the steady-state values of $S$ and $\rho$ are very similar for the two different values of $r_{\rm nuc}$ studied and the dependency on the kinetic parameters remains largely the same. Furthermore, these two quantities are dynamically coupled, as indicated in Figure 2 of the main text as well, with only highly ordered systems achieving large surface densities. The apparently complex dependency of $S$ and $\rho$ on $r_{\rm on}$ and $\tau_{\rm det}$ can in fact be better understood as being controlled only by the intrinsic filament size $N_{\rm c}$ (a measure of the persistence of the treadmilling kinetics) with a clear transition towards order and high density around $N_{\rm c} \sim 100$ monomers (see Figure S10 and Figure 2 in the main text). Importantly, we find that this transition is largely independent, not only of the nucleation rate $r_{\rm nuc}$, but also of structural and dynamical properties of the filaments as changing the stiffness of these ($l_{\rm p}$) or the diffusion coefficient of the monomers ($D$) does not affect the transition much (Figure S10).

Like in single filament simulations, we define the treadmilling velocity as the average displacement of the filament head $\bar{v} = \langle dr_{\rm head}/dt \rangle$, now averaging over the full lifetime of the filament. To measure individual alignment of filaments with the bulk we define the individual nematic order parameter $S_i = 0.5(3\langle(\mathbf{u}_j\mathbf{u}_k)^2\rangle_{\{j,k\}\in i} - 1)$ where we average over all pairs of bond directors $\{j,k\}$ which include a bond belonging to filament $i$. With this measure, $S_i = -0.5$ means the filament is perpendicular to the average collective orientation and $S_i = 1.0$ that the filament is parallel to the bulk of the system. With this measure we can then measure average filament lifetimes and velocities for different degrees of alignment (different values of $S_i$). As shown in Figure 2 of the manuscript and Figure S11, this unveils that aligned filaments in treadmilling systems live longer and move faster than their misaligned counterparts on average, which leads to global alignment and nematic order as misaligned filaments die out over time (see Supplementary Movie 7 for an illustrative example).

We define the relaxation times to steady-state as the time it takes for the nematic order parameter to stabilise. To find it we compare the instantaneous value of $S_t$ measured every 10 seconds over each trajectory with the long time average $S^*$ from those times. We thus define $S^*(t) = \langle S_{t'} \rangle_{\{t' \geq t\}}$ together with its error $\sigma_S(t)$ defined from $\sigma_S^2(t) = \langle S_{t'}^2 - S^*(t) \rangle_{\{t' \geq t\}}$. We can then define the relaxation time $T_{\rm rel}$ as the first time $t$ for which the difference between the instantaneous value and long-time average of the nematic order is smaller than the error: $T_{\rm rel} = \min(t \iff \|S_t - S^*(t)\| < \sigma_S(t))$ (Figure S12a). From such measurements we can extract probability distribution functions for the relaxation times at different nucleation rates, revealing that even for very large $r_{\rm nuc}$ model treadmilling filaments display $T_{\rm rel}$ on the order of minutes (Figure

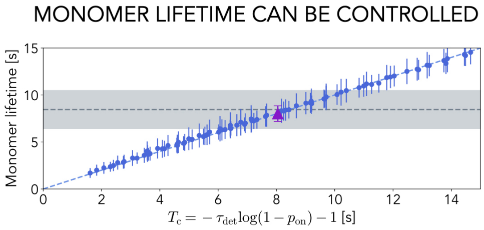


FIG. S5. **Monomer lifetimes** Average monomer lifetimes for different values of $\{r_{on}, \tau_{det}\}$ plotted against the corresponding expected lifetime $T_c$. Each point corresponds to the average over 20 replicas for each parameter set. The dashed line is $y = x$.

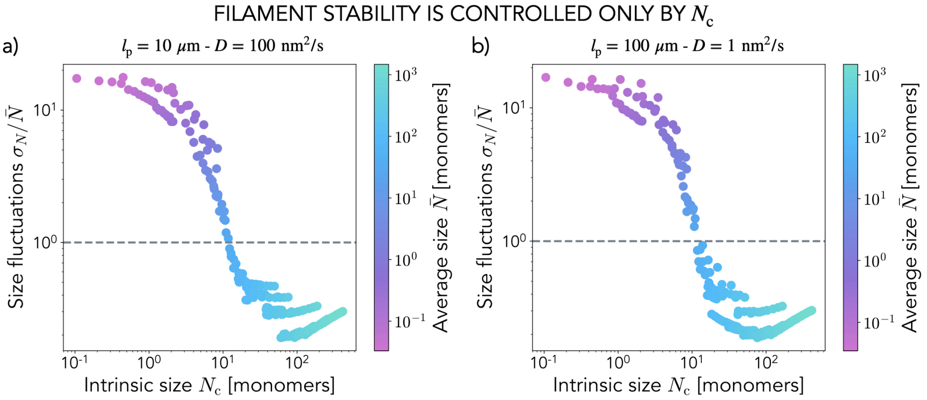


FIG. S6. **Treadmilling stability is controlled by the intrinsic size** Scatter plot of the rescaled filament size fluctuations $\sigma_N/\bar{N}$ against the intrinsic size $N_c = -r_{on}\tau_{det}\log(1-p_{on})$ coloured according to the average filament size $\bar{N}$. As $N_c \gg 10$ monomers the size fluctuations become negligible, indicating a highly stable treadmilling regime.

S12b).

To characterise the locality of the two order parameters $S$ and $P$ we compute these for different perpendicular distances $r_+$ between filaments over the course of a simulation. We define $r_+$ as the distance between monomers $i$ and $j$ along the perpendicular direction to the local filament director at monomer $i$ ($u_i$). We average over $N = 10^6$ pairs of monomers binning them by $r_+$ (bin size $\Delta r_+ = \sigma$) in a single frame of a simulation and consider frames at 1 minute intervals for long times to focus on the steady-state regime. As shown in Figure S13, the polar order decays very quickly as we move away from a filament while the nematic order remains high for large distances. This feature is characteristic of the polar lanes that form in the ordered regime of treadmilling, whereby parallel alignment only spans a few filaments that form a single lane. Note that this behaviour is consistent over time as lanes can grow or shrink but the system never evolves to display global polar order. In Figure S13 we focus on a simulation for $r_{on} = 8$ s$^{-1}$ and $\tau_{det} = 15$ s for the same conditions as in Figure 2 of the main text ($L = 200\sigma$, $l_p = 10\ \mu$m and $D = 100$ nm$^2$/s).

### F. Death by misalignment through treadmilling kinetics drives ordering in dilute conditions as well

In the results presented in the main text for unbiased collective treadmilling dynamics we observe that ordered systems tend to build up their surface density to relatively high values (as much as $\rho \sim 0.6\ \sigma^{-2}$). However, FtsZ numbers in bacterial cells are typically estimated around $N_{FtsZ} \sim 5000$, of which roughly $30-40\%$ are in the Z-ring [19]. For rings roughly $\sim 200$ nm wide in cells 1 $\mu$m in diameter such estimates should limit the surface density to $\rho_{max} \sim 0.08\ \sigma^{-2}$. Consequently, we thought it would be important to test whether the mechanism we describe in this work, whereby filaments collectively organise via the death of misaligned ones, still holds at more *in vivo*-like dilute

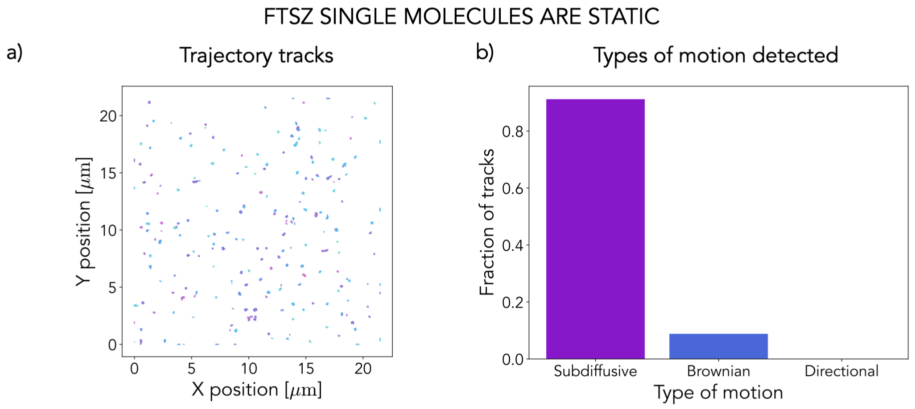


FIG. S7. **Single FtsZ monomers are static a)** Single FtsZ molecules trajectory tracks ($N = 288$). **b)** Types of motion detected (fraction of total tracks) after analysis of the collected tracks ($N = 136$).

conditions. For this purpose we repeated the same simulations as presented in Figure 2 of the main text now constraining our system to a maximum number of particles $N_{\rm max} = 3200$ which, for systems of $L = 200\sigma$, gives $\rho_{\rm max} \sim 0.08\ \sigma^{-2}$. In this case we fix $K_{\rm bend} = 10^4\ k_{\rm B}T$ ($l_{\rm p} = 100\ \mu$m) and $D = 10$ nm$^2$/s. We find that, if we simulate treadmilling kinetics in the ordered region of the parameter space dilute systems still order, reaching large values of the nematic order $S$ while saturating their surface density to $\rho_{\rm max} \sim 0.08\ \sigma^{-2}$ (Figure S14a-b). Here as well we can characterise the dependence of filament lifetimes and velocities on their individual alignment $S_i$, which gives the same results as for the unconstrained simulations: filaments on average live longer and treadmill faster when they are more aligned with their surroundings, a trademark of the mechanism at play (Figure S14c-d). These results thus show that treadmilling filaments need not be at large surface density to order. Instead, the ordering process is driven solely by their kinetics and in turn allows for a surface density buildup if no limits on concentration are set.

#### G. Treadmilling is essential for ordering

We provide further proof of the treadmilling origin of the order transition we observe in our model by artificially arresting the dynamics at different stages along relaxation. We implement this by turning off filament shrinking ($r_{\rm off} = 0$) at different arrest times $t_{\rm arr}$. We perform simulations in a system of size $L = 200\sigma$ for two kinetic parameter sets – one that remains disordered and one that orders ($\{r_{\rm on} = 4\ {\rm s}^{-1}, \tau_{\rm det} = 4\ {\rm s}\}$ and $\{r_{\rm on} = 8\ {\rm s}^{-1},\ \tau_{\rm det} = 15\ {\rm s}\}$) over 10 minutes. We find that in the absence of treadmilling the system freezes and saturates in a disordered manner as a result of turnover inhibition. For systems that remain disordered in steady-state arresting treadmilling only has the effect of freezing the system and saturating the surface density $\rho$, but the nematic order $S$ remains low, albeit slightly larger than for unperturbed treadmilling as thermal fluctuations and collisions can foster local alignment for long-lived filaments (Figure S15a,c). For systems that order in steady-state when their dynamics are unperturbed we find that arresting treadmilling decreases the order as newly nucleated defects are not dissolved. Consequently, we find that the earlier we kill the off rate the more disordered the system becomes as there is more room for defects and that if treadmilling is arrested after high order is reached then the system remains ordered (Figure S15b,c). Here again kinetic arrest results in frozen systems that saturate the surface density $\rho$.

#### H. High-speed AFM data

Raw High-Speed Atomic Force Microscopy (HS-AFM) images (see Section M) were analysed using the OrientationJ plugin for imageJ [20] to obtain vector fields for each frame after adequate thresholding (Figure S16b). Only pixels with high intensity ($I > 122.5$, $I_{\rm max} = 255$) are retained for nematic analysis. We define the surface density from AFM images as $\rho_t = \sum_{I>122.5} {\rm pxl}/N_{\rm pxl}$ where $N_{\rm pxl}$ is the total number of pixels in the frame. Similarly to simulations analysis, we define the nematic order pa-

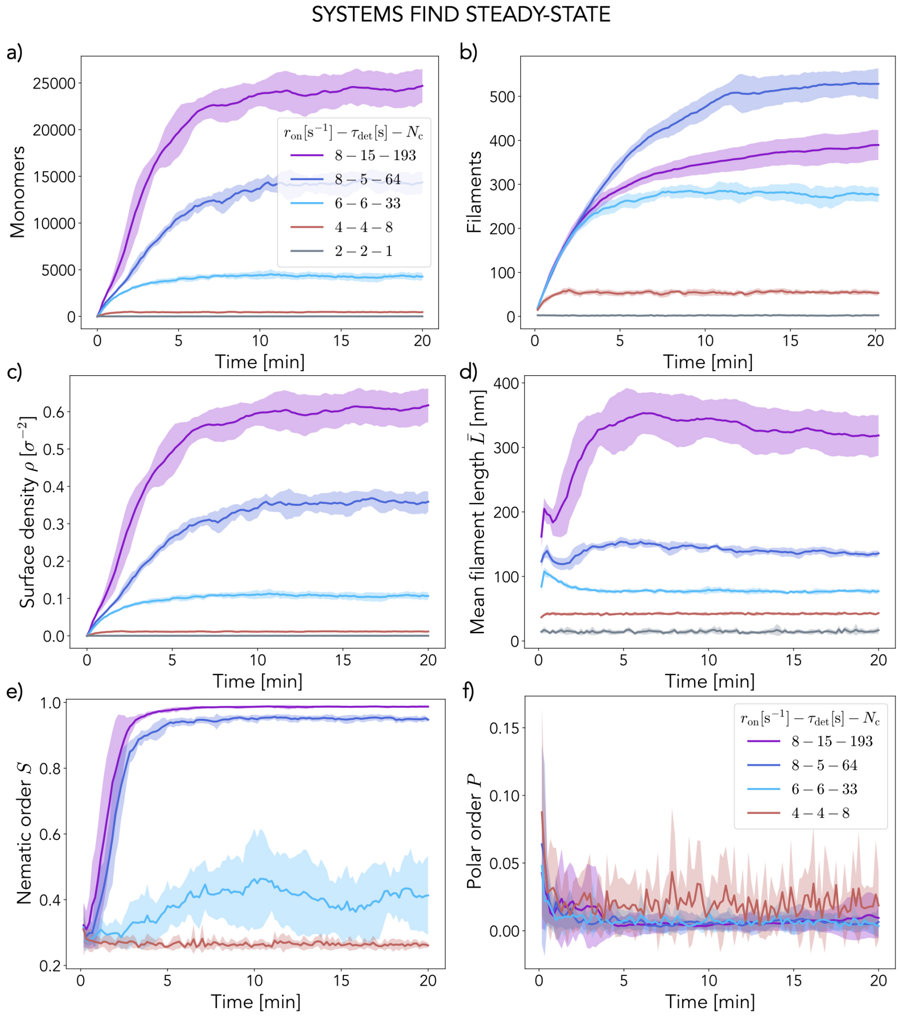


FIG. S8. **Collective steady-state evolution of treadmilling systems** Temporal evolution of different relevant system variables in five representative systems of treadmilling filaments showing stabilisation around constant values after a few minutes. **a)** Total number of monomers. **b)** Total number of filaments. **c)** Surface density of monomers. **d)** Average filament length. **e)** Nematic order $S$. **f)** Polar order $P$. Each curve is the average over $N = 10$ replicas (shaded region is the standard deviation) for different kinetic parameters $\{r_{\rm on}, \tau_{\rm det}\}$ and hence intrinsic size $N_{\rm c}$ (see legend). In all cases $r_{\rm nuc} = 1\ {\rm s}^{-1}$, $L = 200\sigma = 1\ \mu$m, $l_{\rm p} = 10\ \mu$m and $D = 100\ {\rm nm}^2$/s.

rameter as $S_t = \langle 0.5(3(\mathbf{u}_i\mathbf{u}_j)^2 - 1)\rangle_{\{i,j\}}$ where $\{i,j\}$ denotes all pairs of vectors in the high intensity regions. We perform this analysis both on wild-type *E. coli* FtsZ and the L169R mutant that displays inhibited GTPase activity and turnover [21] reconstituted on supported lipid bilayers (see Figure 2 of the manuscript for results).

### I. Implementing the spatio-temporal modulation of kinetics

We incorporate the spatio-temporal modulation of FtsZ polymerisation kinetics by partner proteins and other positioning systems into our model as an instantaneous switch in the growth and nucleation rates at time $t = 0$ while keeping the detachment time $\tau_{\rm det}$ constant. We assume FtsZ polymerisation is initially unperturbed and thus growth and nucleation rates take on a uniform value across the cell

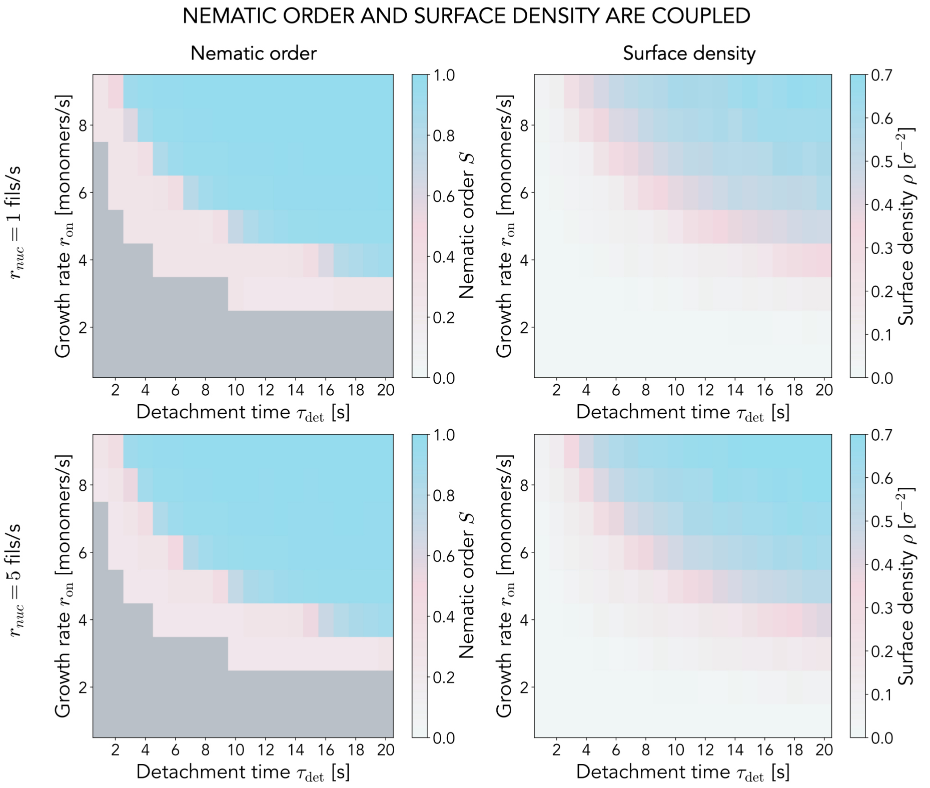


FIG. S9. **Collective filament properties are independent of the nucleation rate** Dependence of the nematic order $S$ and surface density $\rho$ on the two kinetic parameters $\{r_{\text{on}}, \tau_{\text{det}}\}$ for $r_{\text{nuc}} = 1\ \text{s}^{-1}$ (top row) and $r_{\text{nuc}} = 5\ \text{s}^{-1}$ (bottom row). The gray area in the nematic order plots corresponds to non-treadmilling systems that remain essentially empty. Each point corresponds to the average over $N = 10$ replicas in steady state ($t > 10$ minutes out of 20 total) for systems of size $L = 200\sigma$. $l_{\text{p}} = 10\ \mu\text{m}$ and $D = 100\ \text{nm}^2/\text{s}$.

body: $r_{\text{on}}(t \leq 0, Y) = r^0_{\text{on}}\ \text{s}^{-1}$ and $r_{\text{nuc}}(t \leq 0, Y) = r^0_{\text{nuc}}\ \text{s}^{-1}$, where $Y$ is the position on the cell axis. Because chemical patterning systems, condensates and chromosome association partners generally have the combined effect of promoting FtsZ polymerisation around the midcell region while inhibiting it around the poles of the cell [22–32], we model the modulated kinetics for $t > 0$ as a Gaussian distribution of typical width $w_{\text{prof}}$ centered around midcell: $r_{\text{on}}(t > 0, Y) = r^1_{\text{on}}\ \exp(-4Y^2/w^2_{\text{prof}})\ \text{s}^{-1}$ and $r_{\text{nuc}}(t > 0, Y) = r^1_{\text{nuc}}\ \exp(-4Y^2/w^2_{\text{prof}})\ \text{s}^{-1}$, where $r^1_{\text{on}}/r^0_{\text{on}} = r^1_{\text{nuc}}/r^0_{\text{nuc}} > 1$. Note that, since ring widths have been measured experimentally to be around $\sim 100$ nm [33–35], we set $w_{\text{prof}} = 100\ \text{nm} = 20\sigma$ unless stated otherwise. For the data shown in Figure 3 of the manuscript and Figure S18, for instance, we set $r^0_{\text{on}} = 2\ \text{s}^{-1}$, $r^1_{\text{on}} = 8\ \text{s}^{-1}$ and $r^1_{\text{nuc}} = 1\ \text{s}^{-1}$ with $\tau_{\text{det}} = 15$ s, while for the data in Figure 4 and Figure S20 we set $r^0_{\text{on}} = 2\ \text{s}^{-1}$, $r^1_{\text{on}} = 9\ \text{s}^{-1}$ and $r^1_{\text{nuc}} = 1\ \text{s}^{-1}$ with $\tau_{\text{det}} = 15$ s.

To characterise ring condensation in such simulations we quantify monomer surface density distributions along the cell axis over time: $\rho(t, Y) = \sum_i m_i \delta(Y_i - Y)/L$, where $m_i$ represents monomer $i$ and $Y_i$ is its position along the cell axis. $\delta(x)$ is the Dirac delta function. With this we then compute the septal density as the average within the profile width ($\rho_{\text{S}}(t) = \langle \rho(t, \|Y\| \leq w_{\text{prof}}) \rangle$) and the distribution width as the span of the region beyond which the density decays below half the average at midcell. See Figure S17 for an illustration.

#### J. The effect of attractive interactions

To explore the suggested stabilising effects of cross-linking proteins during FtsZ condensation into the Z-ring we introduce a generic Lennard-Jones (LJ) type of attraction between polymers (see Section A).

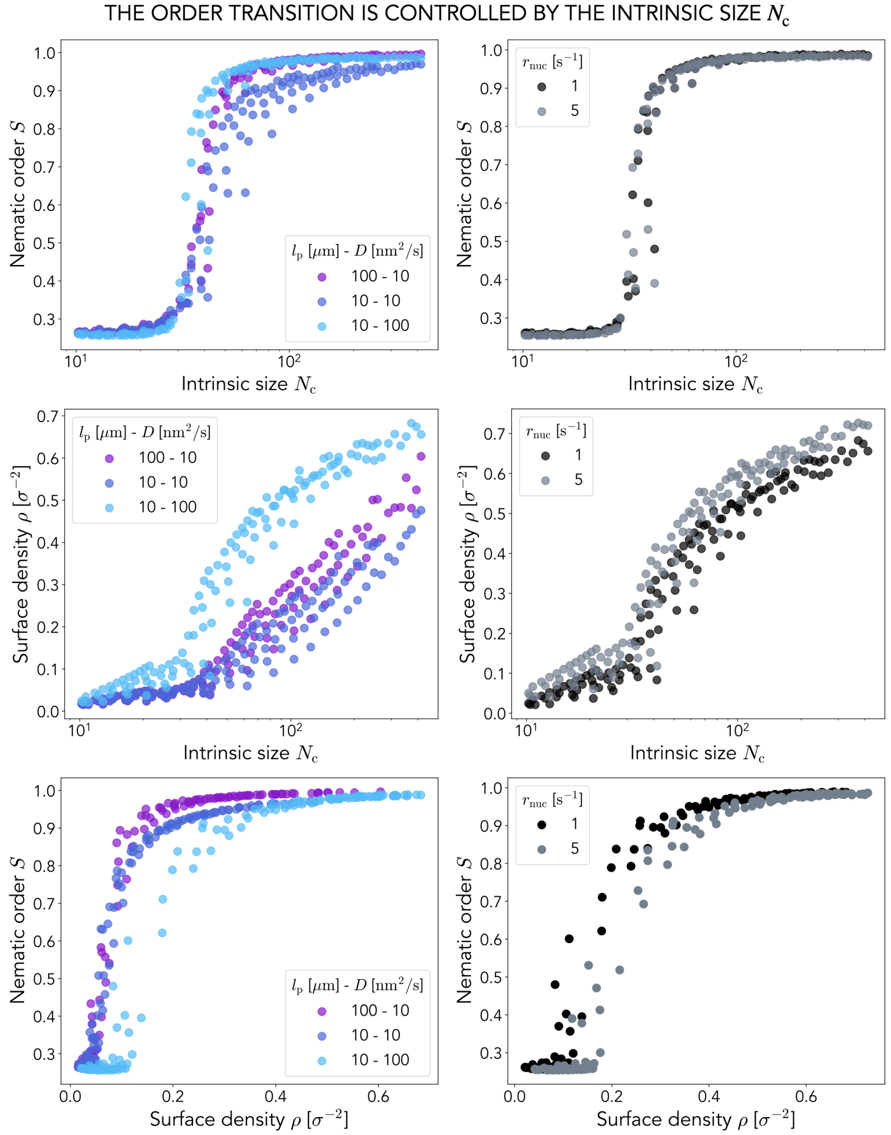


FIG. S10. **Collective order and density are controlled by the intrinsic filament size** Top row: Nematic order $S$ at steady state plotted against the intrinsic filament size $N_c$ for different conditions. Middle row: Surface density $\rho$ at steady state plotted against the intrinsic filament size $N_c$ for different conditions. Bottom row: Coupling of nematic order $S$ and surface density $\rho$ at steady state. Left column: Comparing different filament persistence lengths $l_p$ and monomer diffusion coefficients $D$ (see legend). In all cases $r_{nuc} = 1$ s$^{-1}$. Right column: Comparing different nucleation rates $r_{nuc}$ (see legend). In all cases $l_p = 10$ $\mu$m and $D = 100$ nm$^2$/s. Each point corresponds to the steady state ($t > 10$ minutes out of 20) average over $N = 10$ replicas for each kinetic parameter combination $\{r_{on}, \tau_{det}\} \to N_c$. In all cases $L = 200\sigma$.

To model only cross-linking effects and avoid self-interactions within the filaments we only consider attractive interactions of this type between monomers of different polymers.

As shown in Figure S18 and Figure 3 of the main text, we find that such cross-linking interactions have an important stabilising effect on the ring structures the model produces. We measure the density profile along the cell axis over time and observe that, as expected, cross-linked rings display a

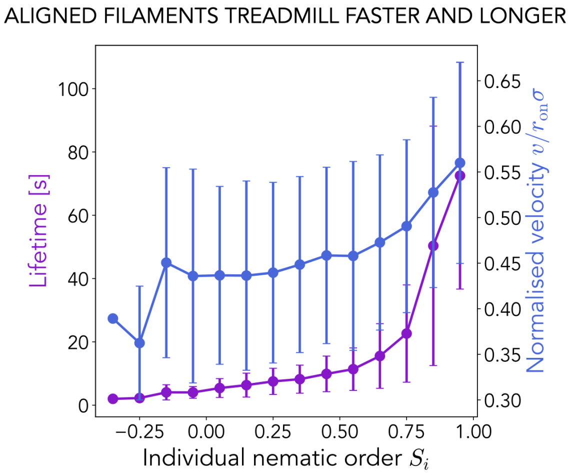


FIG. S11. **Aligned filaments treadmill longer and faster** Filament lifetime and normalised velocity $v/r_{\rm on}\sigma$ for different levels of alignment (characterised by the individual nematic order $S_i$). We consider only simulations for $N_{\rm c} \geq 100$ ($N = 10$ replicas per parameter set), which results in 226077 data points (filaments) in total binned in $S_i$ with width 0.1. We use $r_{\rm on} = 8$ s$^{-1}$, $\tau_{\rm det} = 15$ s and $r_{\rm nuc} = 1$ s$^{-1}$ for $l_{\rm p} = 10$ $\mu$m and $D = 100$ nm$^2$/s in a system of size $L = 200\sigma$.

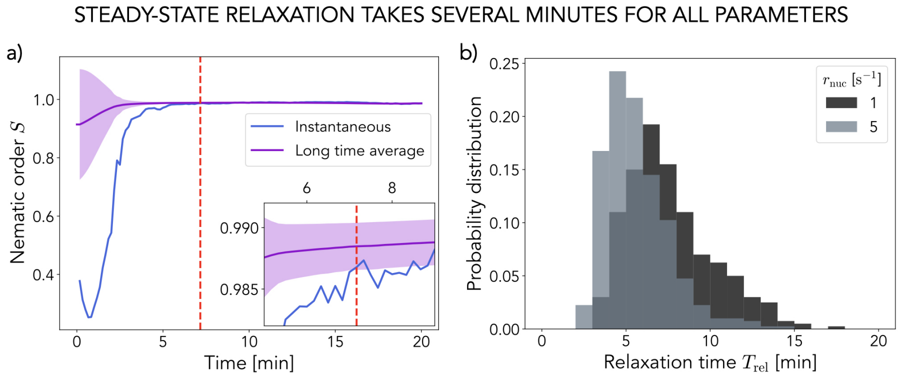


FIG. S12. **Relaxation to steady-state occurs on the order of minutes a)** Example trajectory of the nematic order $S_t$ and its long time average $S^*(t)$ (error $\sigma_S(t)$ is shown by the shaded region) considered to find the relaxation time $T_{\rm rel}$ (dashed vertical line). The inset is a zoom-in around $T_{\rm rel}$. **b)** Probability distribution of relaxation times to nematic order steady-state for two different nucleation rates $r_{\rm nuc}$. The distributions were obtained by running statistics over 40 different parameter sets that order ($N_{\rm c} \geq 100$) and $N = 10$ replicas for each, resulting in 400 data points per curve. In all cases $l_{\rm p} = 10$ $\mu$m, $D = 100$ nm$^2$/s and $L = 200\sigma$.

more condensed configuration, where filaments bundle closer together and most of the monomer population is confined to a narrower stripe around the midcell. As such, the profile displays a narrower and more peaked Gaussian-like distribution around midcell compared to non-cross-linked rings.

#### K. Z-ring dynamics analysis

We perform simulations of model treadmilling filaments subject to geometrical (curvature force $f_{\rm curv} = 5$ $k_{\rm B}T$) and chemical biases (modulation of kinetics at time $t = 0$) as well as cross-linking interactions to explore ring condensation dynamics. To simulate systems in cell-like conditions we set $L = 600\sigma = 3$ $\mu$m such that our systems correspond to cylinders 3 $\mu$m wide and $\sim 1$ $\mu$m in diameter. Fur-

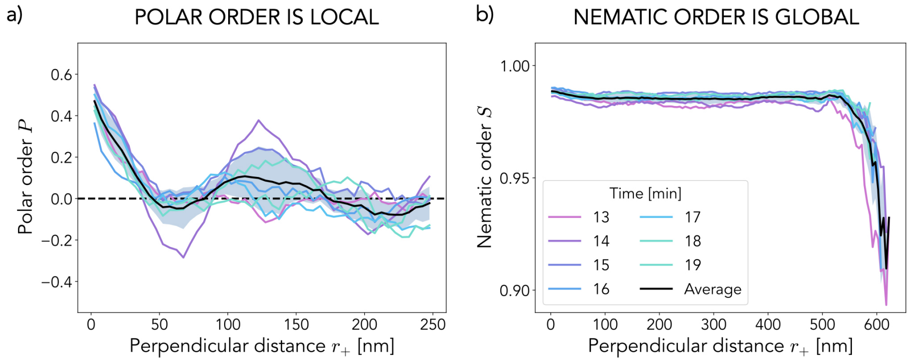


FIG. S13. **Polar order is local while nematic order is global** Polar order parameter $P$ (panel a) and nematic order parameter $S$ (panel b) measured between filaments for different perpendicular distances $r_+$ at several times during the simulation ($r_{\rm on} = 8$ s$^{-1}$, $\tau_{\rm det} = 15$ s, $r_{\rm nuc} = 1$ s$^{-1}$). A consistent sharp decay at small distances emerges for $P$ independent of time while $S$ remains high at large distances. Different colours correspond to different times and the black line represents the average over these (the shaded region indicates the error). For each curve $N = 10^6$ pairs of bonds were sampled in the system ($L = 200\sigma$, $l_{\rm p} = 10$ $\mu$m, $D = 100$ nm$^2$/s).

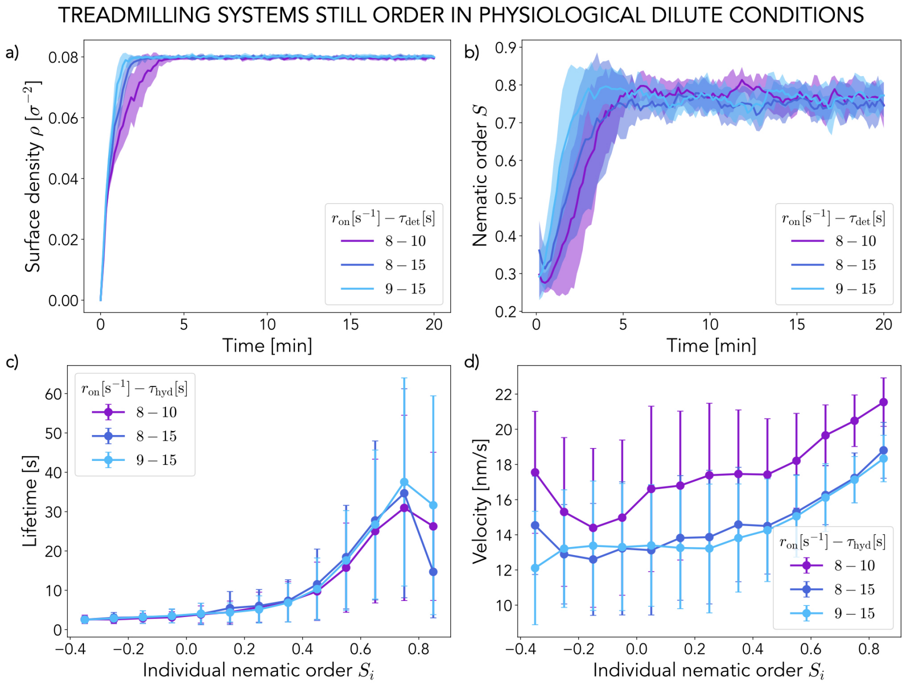


FIG. S14. **Model treadmilling filament systems constrained to a dilute regime still order by misaligned filaments dying out** a-b) Time evolution of the surface density $\rho$ and nematic order $S$ respectively. The system saturates to $\rho_{\rm max} = 0.08$ $\sigma^{-2}$ but still reaches a globally ordered steady state $S_{t\to\infty} \sim 0.8$. Solid lines are the average over $N = 10$ different replicas (thin, faded lines). c-d) Average filament lifetimes and velocities respectively for different levels of individual alignment (characterised by $S_i$). Aligned filaments treadmill faster and display live longer on average. Points are the average for each alignment bin (width 0.1) and error bars correspond to the standard deviation. Statistics over $N = 10$ replicas. Different kinetic parameters are coloured according to the legend. $L = 200\sigma$, $l_{\rm p} = 100$ $\mu$m, $D = 10$ nm$^2$/s.

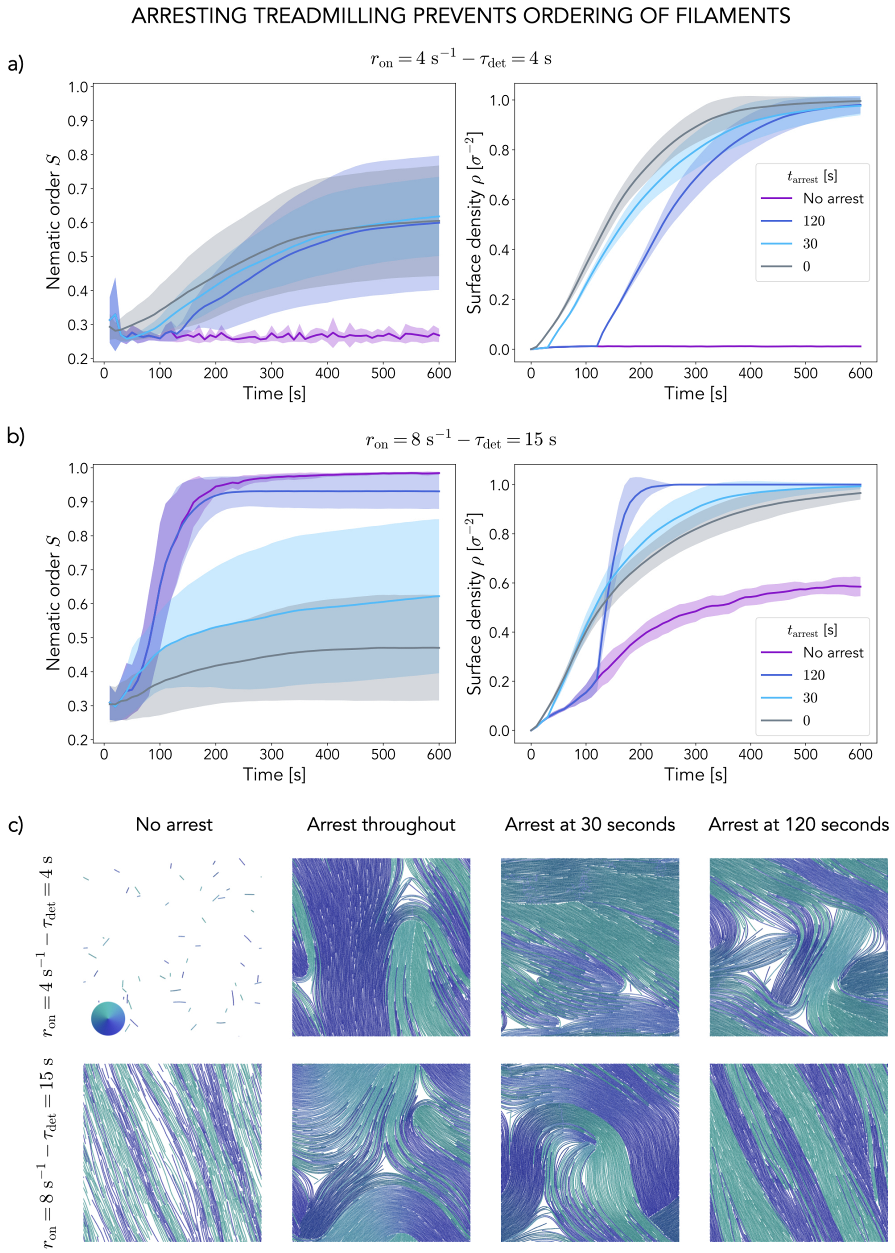


FIG. S15. **Arrested treadmilling leads to disorder a,b)** Surface density $\rho$ and nematic order $S$ curves over time for different scenarios of treadmilling arrest (see legend, $t_{arr}$ is the time when the off rate is killed). We consider two different kinetic parameters, $r_{on} = 4$ s$^{-1}$ and $\tau_{det} = 4$ s (a) and $r_{on} = 8$ s$^{-1}$ and $\tau_{det} = 15$ s (b) normally resulting in disorder and order respectively. Curves correspond to the average over $N = 10$ replicas. **c)** Representative snapshots of the system after 10 minutes for the different kinetic parameters and arrest times. Filaments are coloured according to their orientation (see wheel). In all cases $L = 200\sigma$, $l_p = 10$ $\mu$m and $D = 100$ nm$^2$/s.

thermore, it is known that a total of around $\sim 5000$ FtsZ monomers are present in a typical *B. subtilis* cell, of which an estimated $\sim 30 - 40\%$ are found in the Z-ring [19], so we constrain our systems to a maximum number of particles $N_{max} = 2000$. Finally, because FtsZ/FtsA composites are expected to be quite stiff and present negligible diffusion in experiments [1, 5, 18, 36, 37], we set $l_p = 100$ $\mu$m

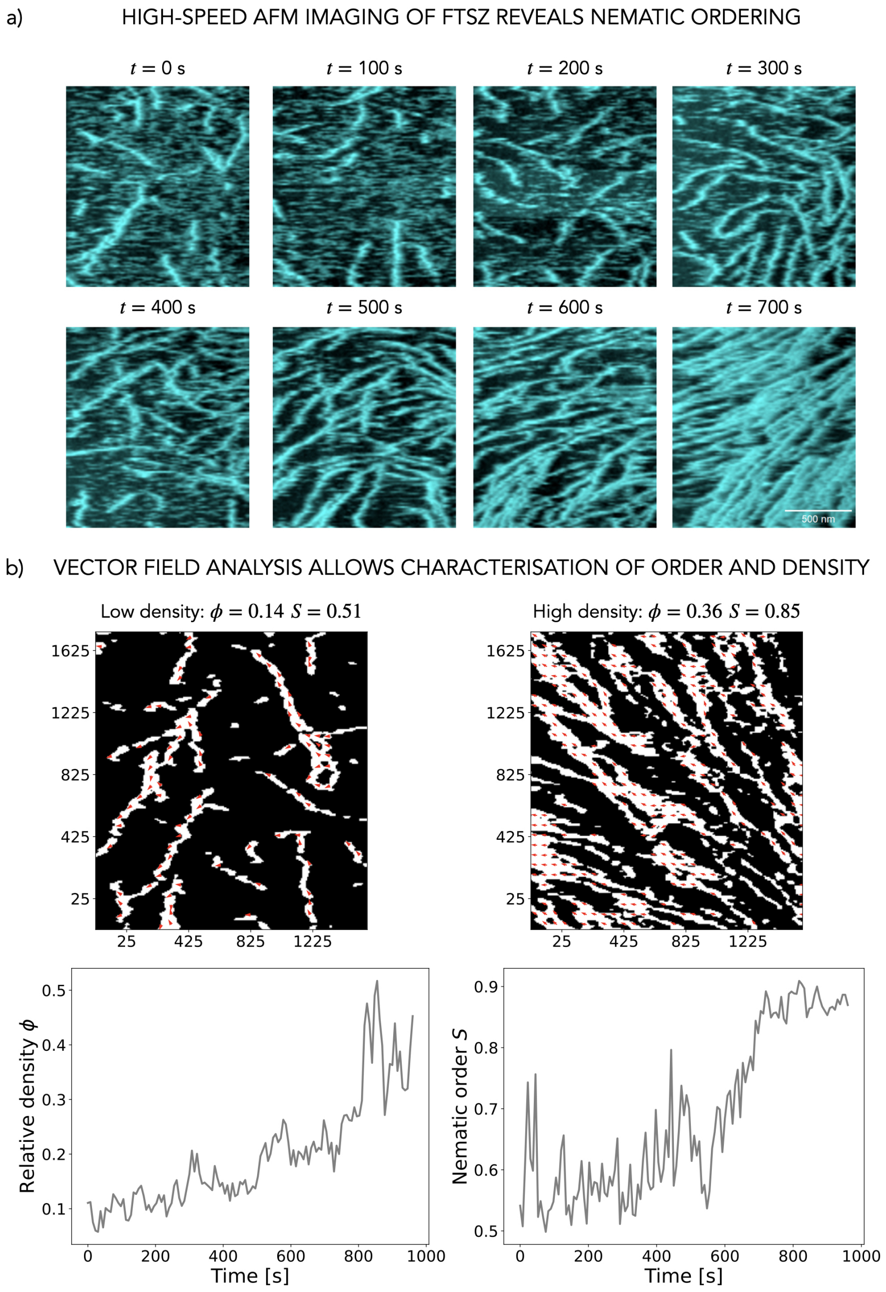


FIG. S16. **AFM imaging of reconstituted FtsZ a)** Time series of a system of wild-type FtsZ from *E. coli* undergoing a nematic order transition when reconstituted on a supported lipid bilayer. Images are acquired by High-Speed Atomic Force Microscopy. Scale bar: 500 nm. **b)** Top row: Vector field analysis of two representative snapshots ($t = 150$ seconds and $t = 900$ seconds respectively) of the system in panel a, including the resulting relative surface density $\rho$ and nematic order $S$. White corresponds to measured filaments and black to empty space. Bottom row: Time series of the relative surface density $\rho$ and nematic order parameter $S$ as measured by vector field analysis of HS-AFM images.

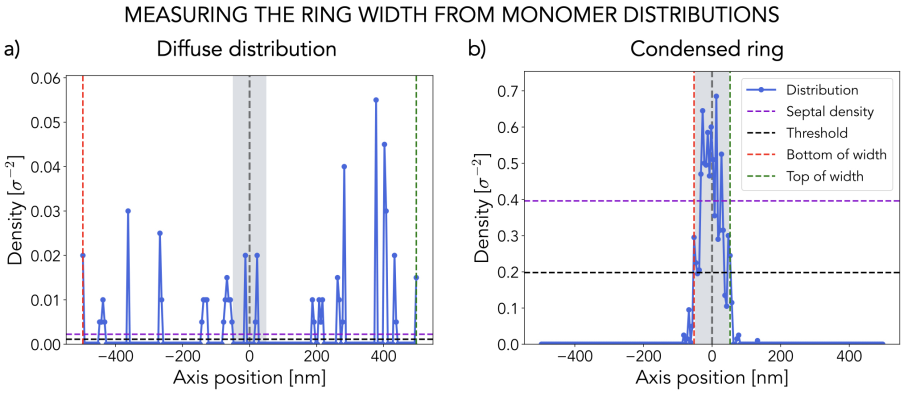


FIG. S17. **Measuring ring density and width** Monomer surface density distribution along the cell axis for two representative simulation frames (diffuse distribution – panel a, and condensed ring – panel b). The blue curve indicates the measured distribution of monomers. The purple dashed line the average density in the septal region ($\|Y\| \leq w_{\rm prof}$, gray shaded region). The black dashed line indicates the established threshold for the distribution width (half of the septal density). The red and green vertical lines indicate the bottom and top edges of the measured width respectively (where the distribution value falls below threshold).

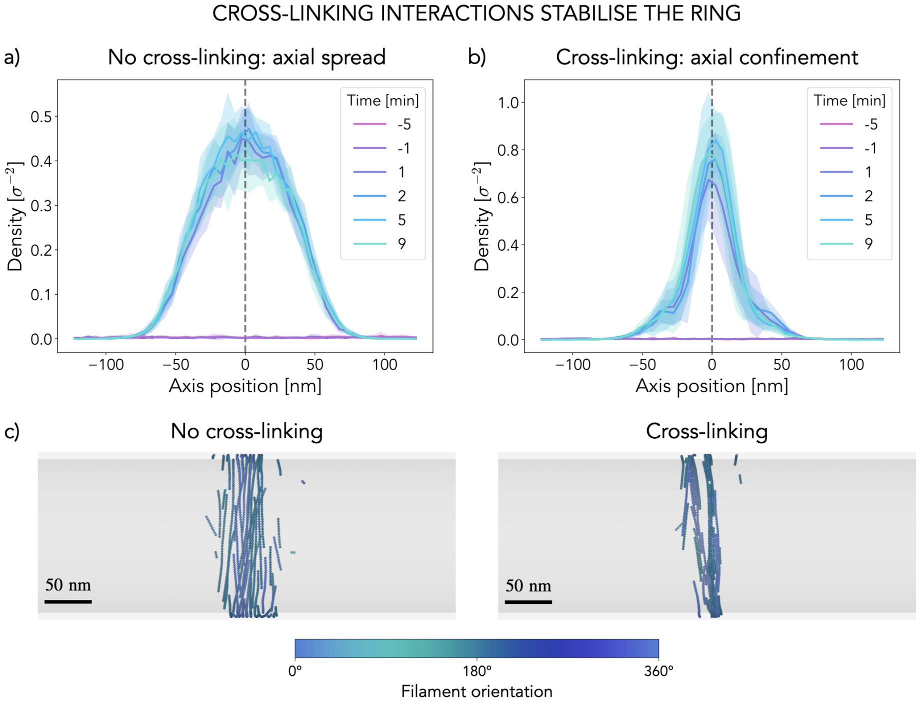


FIG. S18. **Cross-linking interactions stabilise and confine the ring axially** Monomer density profile along the cell axis for non-cross-linked (panel a) and cross-linked (panel b) filaments at different times along the simulation. Note the different scales in the two panels. Each curve is the result of averaging 1 minute around the indicated time ($\pm 30$ s) over $N = 10$ replicas (solid lines are the average and shaded regions the standard deviation). In all cases $L = 200\sigma = 1\mu$m, $l_{\rm p} = 10\ \mu$m and $D = 100$ nm$^2$/s. To simulate ring formation we impose a kinetics modulation with parameters $r^0_{\rm on} = 2$ s$^{-1}$, $r^1_{\rm on} = 8$ s$^{-1}$ and $r^1_{\rm nuc} = 1$ s$^{-1}$ for $\tau_{\rm det} = 15$ s and $w_{\rm prof} = 100$ nm.

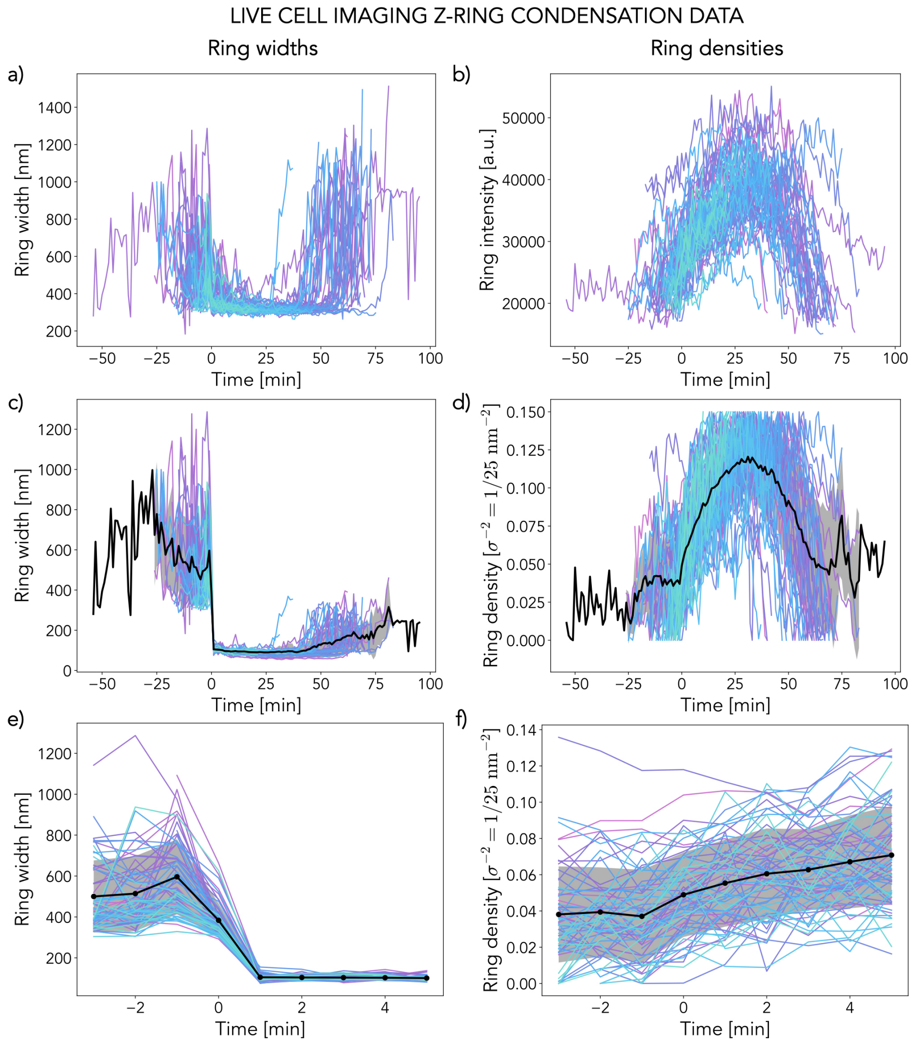


FIG. S19. **Experimental *in vivo* Z-ring data a)** Raw ring width trajectories over time aligned for width collapse at $t = 0$ minutes. **b)** Raw ring intensity trajectories over time aligned for width collapse at $t = 0$ minutes. **c)** Ring width trajectories for live cells adjusted for diffraction limits (estimated final ring width is 100 nm). **d)** Rescaled ring intensity trajectories to match estimated densities of up to 0.15 $\sigma^{-2} = 0.006$ nm$^{-2}$ **e)** Ring width trajectories for live cells adjusted for diffraction limits around the collapse at time $t = 0$ (estimated final ring width is 100 nm). **f)** Rescaled ring intensity trajectories around $t = 0$. In all cases the black curve and shaded region correspond to the average and standard deviation across all individual cells ($N = 67$, different colours).

and $D = 1$ nm$^2$/s. For kinetics modulation we impose a transition in growth and nucleation rates following $r^0_{\rm on} = 2$ s$^{-1}$, $r^1_{\rm on} = 9$ s$^{-1}$ and $r^1_{\rm nuc} = 1$ s$^{-1}$. In all cases we keep $\tau_{\rm det} = 15$ s. As for the width of the profile, because Z-rings have been measured to be around 100 nm wide for *Caulobacter* [33], *E. coli* [34] and especially *B. subtilis* [35] (our species of interest) we set $w_{\rm prof} = 100$ nm. Note as well that because FtsZ seems confined to an area of around $\sim 500$ nm around midcell in early stages of the cell cycle we also impose a Gaussian kinetics profile for $t < 0$: $r_{\rm on}(t \leq 0, Y) = r^0_{\rm on}\,\exp(-4Y^2/w^2_{\rm conf})$ s$^{-1}$ and $r_{\rm nuc}(t \leq 0, Y) = r^0_{\rm nuc}\,\exp(-4Y^2/w^2_{\rm conf})$ s$^{-1}$, where $w_{\rm conf} = 500$ nm. We simulate our rings over time $t \in [-10, 10]$ minutes and measure distribution widths along the cell axis together with average surface densities in the septal region ($|Y| \leq w_{\rm prof}$) over time.

**Rescaling of *in vivo* data** We perform high-resolution imaging of FtsZ filament dynamics in live *Bacillus subtilis* cells throughout all stages of division. For this purpose we developed a custom microscopy setup based on vertical cell immobilisation by nanostructures termed VerCINI [38]. With this approach we obtain quantitative measurements of Z-ring widths and fluorescence intensities over time for $N = 67$ cells [39] which we align in time such that all rings nucleate at $t = 0$ minutes (Figure S19a-b). In this study we focus on the dynamics of Z-ring formation. In particular, we look at ring nucleation, characterised by the sudden ring width collapse shortly after $t = 0$, which we observe consistently across all imaged cells (Figure S19a,c,e). We also look at ring maturation, characterised by a sustained FtsZ intensity increase in the septal region over time after ring nucleation, again consistently displayed across all imaged cells (Figure S19b,d,f). However, raw measurements of ring widths and FtsZ intensities cannot be used for quantitative comparison with simulations of model treadmilling filaments. We thus work with rescaled quantities. For experimental measurements of Z-ring widths, which are diffraction limited and do not capture the final ring width properly, we rescale each individual trajectory such that the long-time average fits superresolution measurements of 100 nm [33–35] without affecting the widths before condensation ($t < 0$). For FtsZ intensity in the septal region, which we take as a proxy for monomer density, we rescale each individual trajectory such that its values are confined between $0.000 - 0.006$ nm$^{-2}$ = $0.00 - 0.15$ 1/25 nm$^{-2}$ = $0.00 - 0.15$ $\sigma^{-2}$, the estimated minimum and maximum FtsZ surface densities in live Z-rings. These estimates are based on a total number of 2000 FtsZ molecules [19] confined to a cylindrical region $\sim 100$ nm wide and $\sim 1$ $\mu$m in diameter [33–35]). In this way we obtain rescaled trajectories for ring width and density which are then directly comparable to simulation results (Figure 4).

Experimental results of in vivo FtsZ dynamics presented here are re-analyses of raw data first presented in [39] - full details of sample and strain preparation, data acquistion and analysis methods, and raw experimental data can be found in that study.

**Resampling of simulation data** Because the data acquisition is different in simulations and *in vivo* experiments simulation measurements need to be resampled for proper quantitative comparison with *in vivo* data. The frame rate in our simulations is 10 seconds (Figure S20a-b) but experiments produce 1 data point per minute (Figure S19) so for proper comparison we average simulation data over $1-$minute intervals across all replicas ($N = 10$) such that a single point and error value are produced for each minute of simulation (Figure S20c-d). Note that independently of the time sampling method average trajectories always display an abrupt rapid distribution collapse around $t = 0$ (Figure S20a) and a slow sustained accumulation of monomers to the septal region (Figure S20b).

### L. Arrested treadmilling stops condensation in simulations

To test how much the ring condensation observed in simulations is dependent on treadmilling dynamics we perform additional simulations of the same system (with kinetic parameters $r^0_{\text{on}} = 2$ s$^{-1}$, $r^1_{\text{on}} = 9$ s$^{-1}$, $r^1_{\text{nuc}} = 1$ s$^{-1}$, $\tau_{\text{det}} = 15$ seconds and profile width $w_{\text{prof}} = 100$ nm) but now arrest treadmilling at different stages of ring condensation and maturation. We arrest tredmilling by turning off the depolymerisation reaction, such that $p_{\text{off}} = 0$ after a certain time $t_{\text{arrest}}$. This has the effect of stopping monomer turnover. Like for the *in vivo* data comparison, we work with systems of size $L = 600\sigma = 3$ $\mu$m with a maximum amount of monomers $N_{\text{max}} = 2000$ and set $l_{\text{p}} = 100$ $\mu$m and $D = 1$ nm$^2$/s. We find that if we turn off the dynamics before the onset of condensation ($t = 0$) the filament population fails to localise to the midcell region (Figure S21) and only the existing structures grow until the maximum number of monomers is reached. Indeed, instead of ring-like dynamic structures we now obtain frozen long filaments that remain disperse along the cell axis (Figure S21b). Note that if we inhibit treadmilling during maturation the ring instead remains condensed but becomes frozen in arbitrary configurations (Figure S21b), which might have important consequences for the recruitment of downstream divisome proteins.

### M. Reconstitution of FtsZ *in vitro*: Materials and Methods

**Protein biochemistry** Proteins used in this study, FtsZ and FtsA, were purified as previously described [2].

**Preparation of coverslips** We used piranha solution (30% $H_2O_2$ mixed with concentrated $H_2SO_4$ at a $1:3$ ratio) to clean the glass coverslips for 60 min. This was followed by extensive washes with double-distilled $H_2O$, 10 min sonication in dd$H_2O$

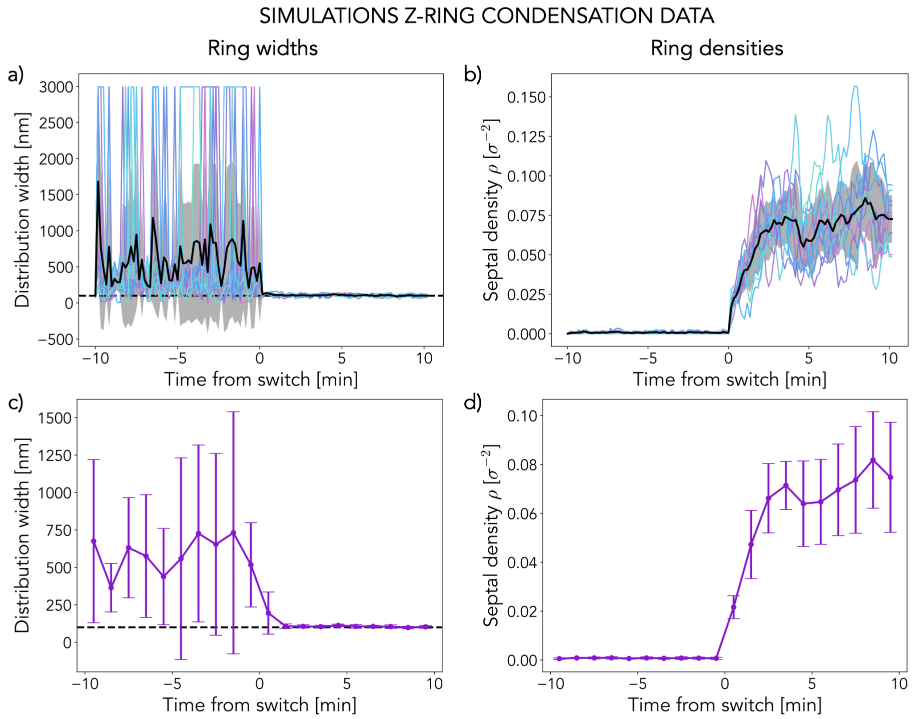


FIG. S20. **Simulation Z-ring data a)** Raw ring width trajectories over time. 1 point every 10 seconds. Black line is the average across $N = 10$ replicas. **b)** Raw ring density trajectories over time. 1 point every 10 seconds. Black line is the average across $N = 10$ replicas. **c)** Time-averaged ring width trajectories over time, 1 data point per minute. Each data point corresponds to the average of all values in the 1 minute interval across all $N = 10$ replicas. Error bars are the standard deviation. **d)** Time-averaged ring density trajectories over time, 1 data point per minute. Each data point corresponds to the average of all values in the 1 minute interval across all $N = 10$ replicas. Error bars are the standard deviation. Simulations were performed for $r^0_{\text{on}} = 2$ s$^{-1}$, $r^1_{\text{on}} = 9$ s$^{-1}$, $r^1_{\text{nuc}} = 1$ s$^{-1}$, $\tau_{\text{det}} = 15$ s and $w_{\text{prof}} = 100$ nm. Systems are $L = 600\sigma = 3$ $\mu$m, $l_{\text{p}} = 100$ nm and $D = 1$ nm$^2$/s.

and again washing in dd$H_2O$. The coverslips were used within one week and were stored in dd$H_2O$ water. Furthermore, before coverslips were used to form supported lipid bilayers, they were dried with compressed air and treated for 10 min with a Zepto plasma cleaner (Diener electronics) at maximum power. As reaction chambers we used 0.5 ml Eppendorf tubes missing the conical end, which were glued on the coverslips with UV glue (Norland Optical Adhesive 63) and exposed to ultraviolet ligth for 10 min.

**Preparation of small unilamellar vesicles (SUVs)** DOPC (1,2-dioleoyl-sn-glycero-3-phosphocholine) and DOPG (1,2-dioleoyl-sn-glycero-3-phospho-(1'-rac-glycerol)), which were purchased from Avanti Polar Lipids, at a ratio of 67 : 33 mol% were used. The lipids in chloroform solution were mixed inside a glass vial in the appropriate volumes and dried with filtered $N_2$ for a thin lipid film. Remaining solvent was removed by putting the lipids in a vacuum desiccator for 2 h. Afterwards swelling buffer (50mM Tris-HCl [pH 7.4] and 300mM KCl) was added to the lipid film to obtain a lipid concentration of 5mM. After incubating the suspension for 30 min at room temperature, the multilamellar vesicles were vortexed rigorously and freeze–thawed (8×) in dry ice or liquid $N_2$. The liposomes were tip-sonicated using a Q700 Sonicator equipped with a 0.5mm tip (amplitude = 1, 1 second on, 4 seconds off) for 25 min on ice to obtain SUVs. Finally, the vesicles were centrifuged for 5 min at 10,000g and the supernatant was stored at 4°C in an Argon atmosphere and used within one week.

**Preparation of supported lipid bilayers (SLB) for TIRF** SLBs were prepared by diluting the

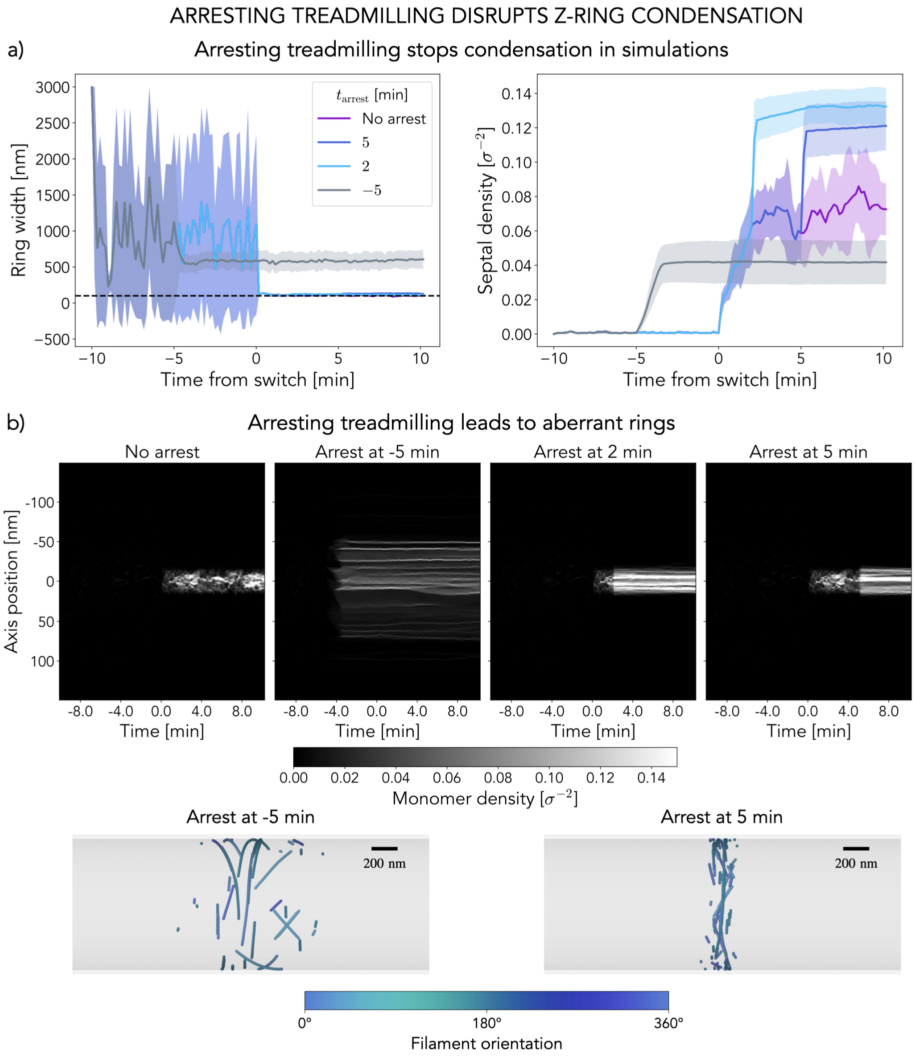


FIG. S21. **Arrested treadmilling stops condensation in simulations a)** Ring width and density curves over time for normal condensation simulations and arrested treadmilling at different times (see legend). Curves correspond to the average over $N = 10$ replicas and shaded regions to the standard deviation. The dashed line in the left panel indicates the target width of 100 nm. **b)** Representative examples of the evolution in time of the monomer distribution along the axis of the cell for four different treadmilling arrest conditions. Even after condensation, treadmilling inhibition severely affects ring structure. Two representative snapshots of the final configuration for treadmilling arrest before and after condensation are shown. Note that monomers in simulations are rendered with size 20 nm instead of the actual 5 nm for visualisation purposes, scale bar is 200 nm. In all cases $L = 600\sigma$, $l_{\mathrm{p}} = 10\ \mu$m, $D = 1$ nm$^2$/s for $r^0_{\mathrm{on}} = 2\ \mathrm{s}^{-1}$, $r^1_{\mathrm{on}} = 9\ \mathrm{s}^{-1}$, $r^1_{\mathrm{nuc}} = 1\ \mathrm{s}^{-1}$, $\tau_{\mathrm{det}} = 15$ s and $w_{\mathrm{prof}} = 100$ nm.

SUV suspension to a concentration of 0.5mM with reaction buffer (50mM Tris-HCl [pH 7.4], 150mM KCl and 5mM $MgCl_2$) supplemented with 5mM $CaCl_2$. SLBs were incubated for 30 min at 37°C and non-fused vesicles were washed away by $8 \times 200\mu$L washes with reaction buffer. The membranes were used within 4 hours.

**Total internal reflection fluorescence (TIRF) microscopy** Experiments were performed using an iMIC TILL Photonics microscope equipped with a 100× Olympus TIRF NA 1.49 differential interference contrast objective. The fluorophores were excited using laser lines at 488 or 640 nm. The emitted fluorescence from the sample was filtered using an Andromeda quad-band bandpass filter (FF01-446-523-600-677). For the dual-colour experiments, an Andor TuCam beam splitter equipped with a spectral long pass of 640 nm and band pass filter combinations of 525/50 and 679/41 nm were used. Time series were recorded using iXon Ultra 897 EMCCD Andor cameras (X-8499 and X-8533) operating at a frequency of 5 Hz.

**High speed atomic force microscopy (HS-AFM)** A laboratory-built tapping mode (2 nm free amplitude, $\sim 2.2$ MHz) high-speed atomic force microscope (HS-AFM) equipped with a wide-range scanner ($6\mu M \times 6\mu M$) was used to visualize the dynamics of the system. BL-AC10DS-A2 (Olympus) cantilevers were used as HS-AFM scanning probes. The cantilever has a spring constant ($k$) of 0.1N/m and a resonance frequency (f) of 0.6MHz in water or 1.5MHz in air. The dimensions of the cantilever are: 9$\mu$m (length), 2$\mu$m (width), and 0.13$\mu$m (thickness). To achieve high imaging resolution, a sharpened and long carbon tip with low apical radius was made on the existing tip of the cantilever using electron-beam deposition (EBD) as described previously [40–42]. Scanning speed varied from 0.2 to 5 seconds per frame. The number of pixels acquired were adjusted for every measurement depending on the scan size (min: 2nm, max: about 50nm). The in-house designed program "Kodec" was used to read the data generated by HS-AFM. The software stores all parameters, calibration and description given during the measurement and allows to load a whole folder or several movies.

**FtsZ TIRF experiments on SLBs** To visualize treadmilling FtsZ filaments on supported lipid bilayers, we used 0.2$\mu$M FtsA and 1.25$\mu$M Alexa488-FtsZ ($1:4$ mixed with unlabelled FtsZ) in 100$\mu$L of reaction buffer. Additionally, the reaction chamber contained 4mM ATP/GTP and a scavenging system to minimize photobleaching effects: 30mM d-glucose, 0.050mg/ml Glucose Oxidase, 0.016mg/ml Catalase, 1mM DTT and 1mM Trolox. Prior addition of all components a corresponding buffer volume was removed from the chamber to obtain a total reaction volume of 100$\mu$L. The FtsZ filaments were imaged by TIRF at 1 frame every 2 seconds and 50ms exposure time.

Single-molecule experiments were performed as described previously [43]. In short, individual FtsZ proteins were imaged at single-molecule level by adding small amounts of Cy5-labelled FtsZ (100pM) to a chamber with 0.2$\mu$M FtsA and 1.25$\mu$M Alexa488-FtsZ.

**Image processing and analysis** For data analysis, the movies were imported to the FIJI software [44]. For data analysis, raw, unprocessed time-lapse videos were used.

**Treadmilling and autocorrelation analysis of FtsZ filaments and single molecules** Treadmilling dynamics as well as the directional autocorrelation were quantified using an automated image analysis protocol previously developed by our group [45].

**Single-molecule analysis of FtsZ** Single molecules of FtsZ were tracked using the TrackMate plugin from ImageJ [46]. To obtain the residence time of FtsZ, we performed a residence time analysis as described before [43, 47]. Shortly, single molecules were imaged at different acquisition rates (0.1˘2 seconds) and the lifetime of the molecules was extracted from each data set. To account for photobleaching effect, the obtained lifetimes were plotted against the acquisition rate and a linear regression was fitted to this data. The photobleach corrected lifetime was obtained by taking the inverse of the slope of the linear regression.

**Preparation of SLBs for HS-AFM** SUVs were prepared as described above. An ultra-flat muscovite mica layers (1.5 mm diameter) substrate was mounted on a glass stage using a standard 2-component glue. The glass stage was then attached to the scanner with a thin film of nail polish. A drop of acetone was deposited on the stage/scanner interface to ensure a flat nail polish layer. The mounted stage was dried at RT for about 30 min. A fresh cleaved mica layer was used as substrate to form a supported lipid bilayer (SLB) by depositing $\sim 4$ $\mu$L of a mix of 1 mM SUVs suspension in reaction buffer with additional 5 mM $CaCl_2$. To avoid drop breakage, the scanner was flipped upside down and inserted in a custom-made mini-chamber with a thin water film at the bottom (a 500 $\mu$L tube cut on the bottom and glued to a petri dish). The drop was incubated on the stage for at least 30 min. After, the drop was exchanged 5-10 times with 5 $\mu$L of fresh reaction buffer. The stage was immediately inserted in the HS-AFM chamber containing about 80 $\mu$L of the same reaction buffer.

Prior to the addition of the proteins, HS-AFM imaging and indentation were performed to assess the quality of the SLB. When the force-distance curve showed the typical lipid bilayer indentation profile ($\sim 2-4$ nm) the SLB was used in the next steps.

**FtsZ HS-AFM experiments on SLBs** The proteins were added to the chamber with ATP/GTP (4 mM each) and DTT (1 mM). Movies used for the analysis contained $1-2$ $\mu$M for FtsZ and 0.5 $\mu$M FtsA.

**Computing the type of motion of FtsZ single molecules** In this study, we tracked FtsZ single molecules using the TrackMate plugin in ImageJ, and exported the resulting trajectories as xml files. The type of movement exhibited by each trajectory was determined through the use of custom Python code available at `https://github.com/paulocaldas/trajectory_analysis_v2`. Specifically, we fitted a linear equation $y = D\ t^{\alpha}$ to the mean square displacement (MSD) of each FtsZ trajectory, with $\alpha$ representing the scaling exponent used to classify the type of motion. Trajectories with $\alpha < 0.6$ were classified as confined, those with $0.6 < \alpha < 1.2$ as Brownian, and those with $\alpha > 1.2$ as directed. To ensure that no short events displaying directed motion were missed, we performed this analysis on sub-segments of each trajectory (window size = 10 frames). With this approach we count the number of tracks for which a certain type of motion was observed, such that if a track displays both Brownian and confined motion, for instance, then both types of motion would increase their count by +1. The results of this analysis are displayed in Figure S7, which clearly indicates that no directional motion of FtsZ was observed, most of it corresponding to confined motion.

### N. Supplementary Movies

**Supplementary Movie 1**: Single treadmilling filament evolving over 30 s in a box of size $L = 100\sigma$. Here $l_{\mathrm{p}} = 10$ $\mu$m and $D = 100$ nm$^2$/s. Bar scale is 100 nm and the video includes a timestamp.

**Supplementary Movie 2**: Disordered treadmilling system evolution over 20 minutes in a box of size $L = 200\sigma = 1$ $\mu$m. Filaments are nucleated at a rate $r_{\mathrm{nuc}} = 1$ s$^{-1}$ and treadmilling kinetics are set by $r_{\mathrm{on}} = 4$ s$^{-1}$ and $\tau_{\mathrm{det}} = 6$ s, corresponding to the pink region in Figure 2a. Here $l_{\mathrm{p}} = 10$ $\mu$m and $D = 100$ nm$^2$/s. Bar scale is 200 nm and the video includes a timestamp.

**Supplementary Movie 3**: Ordering treadmilling system evolution over 20 minutes in a box of size $L = 200\sigma = 1$ $\mu$m. Filaments are nucleated at a rate $r_{\mathrm{nuc}} = 1$ s$^{-1}$ and treadmilling kinetics are set by $r_{\mathrm{on}} = 8$ s$^{-1}$ and $\tau_{\mathrm{det}} = 15$ s, corresponding to the blue region in Figure 2a. Here $l_{\mathrm{p}} = 10$ $\mu$m and $D = 100$ nm$^2$/s. Bar scale is 200 nm and the video includes a timestamp.

**Supplementary Movie 4**: Arrested treadmilling system evolution over 20 minutes in a box of size $L = 200\sigma = 1$ $\mu$m. Filaments are nucleated at a rate $r_{\mathrm{nuc}} = 1$ s$^{-1}$ and treadmilling kinetics are set by $r_{\mathrm{on}} = 8$ s$^{-1}$ and $\tau_{\mathrm{det}} = 15$ s, but $p_{\mathrm{off}} = 0$ throughout (arrested treadmilling). Here $l_{\mathrm{p}} = 10$ $\mu$m and $D = 100$ nm$^2$/s. Bar scale is 200 nm and the video includes a timestamp. The absence of turnover prevents nematic defect dissolution.

**Supplementary Movie 5**: High-Speed AFM image sequence of *E. coli* FtsZ reconstituted on a supported lipid bilayer, imaged over $\sim 13$ minutes. Bar scale is 500 nm and the video includes a timestamp.

**Supplementary Movie 6**: High-Speed AFM image sequence of mutant L169R FtsZ reconstituted on a supported lipid bilayer, imaged over $\sim 13$ minutes. Bar scale is 500 nm and the video includes a timestamp. The inhibited depolymerisation and turnover prevents nematic defect dissolution.

**Supplementary Movie 7**: Trapped filament dissolution. Simulation trajectory example of a single treadmilling filament (highlighted in purple) colliding and getting trapped against its neighbours until eventual dissolution, as its tail keeps shrinking. Kinetic parameters used: $r_{\mathrm{nuc}} = 1$ s$^{-1}$, $r_{\mathrm{on}} = 8$ s$^{-1}$ and $\tau_{\mathrm{det}} = 15$ s. System is $L = 200\sigma = 1$ $\mu$m in size and $l_{\mathrm{p}} = 10$ $\mu$m and $D = 100$ nm$^2$/s. Bar scale is 100 nm and the video includes a timestamp.

**Supplementary Movie 8**: Three-dimensional reconstruction of a ring condensation trajectory from simulations for *in vivo* conditions. Kinetic parameters: $r^0_{\mathrm{on}} = 2$ s$^{-1}$, $r^1_{\mathrm{on}} = 9$ s$^{-1}$ and $r^1_{\mathrm{nuc}} = 1$ s$^{-1}$ for $\tau_{\mathrm{det}} = 15$ s and $w_{\mathrm{prof}} = 100$ nm. System is $L = 600\sigma = 3$ $\mu$m in size (so $R \sim 1$ $\mu$m) and $l_{\mathrm{p}} = 100$ $\mu$m and $D = 1$ nm$^2$/s. Monomers are rendered with size 20 nm instead of the actual 5 nm for visualisation purposes and are coloured according to orientation (see wheel), scale bar is 200 nm. The video includes a timestamp for which $t = 0$ corresponds to the onset of the chemical pattern.

**Supplementary Movie 9**: Three-dimensional reconstruction of a ring condensation trajectory from simulations for *in vivo* conditions where kinetics are arrested ($p_{\mathrm{off}} = 0$) 5 minutes before the onset of modulation. Kinetic parameters: $r^0_{\mathrm{on}} = 2$ s$^{-1}$,

$r_{\mathrm{on}}^{1} = 9\ \mathrm{s}^{-1}$ and $r_{\mathrm{nuc}}^{1} = 1\ \mathrm{s}^{-1}$ for $\tau_{\mathrm{det}} = 15$ s and $w_{\mathrm{prof}} = 100$ nm. System is $L = 600\sigma = 3\ \mu$m in size (so $R \sim 1\ \mu$m) and $l_{\mathrm{p}} = 100\ \mu$m and $D = 1\ \mathrm{nm}^2/\mathrm{s}$. Monomers are rendered with size 20 nm instead of the actual 5 nm for visualisation purposes and are coloured according to orientation (see wheel), scale bar is 200 nm. The video includes a timestamp for which $t = 0$ corresponds to the onset of the chemical pattern.

**Supplementary Movie 10**: Three-dimensional reconstruction of a ring condensation trajectory from simulations for *in vivo* conditions where kinetics are arrested ($p_{\mathrm{off}} = 0$) 5 minutes after the onset of modulation. Kinetic parameters: $r_{\mathrm{on}}^{0} = 2\ \mathrm{s}^{-1}$, $r_{\mathrm{on}}^{1} = 9\ \mathrm{s}^{-1}$ and $r_{\mathrm{nuc}}^{1} = 1\ \mathrm{s}^{-1}$ for $\tau_{\mathrm{det}} = 15$ s and $w_{\mathrm{prof}} = 100$ nm. System is $L = 600\sigma = 3\ \mu$m in size (so $R \sim 1\ \mu$m) and $l_{\mathrm{p}} = 100\ \mu$m and $D = 1\ \mathrm{nm}^2/\mathrm{s}$. Monomers are rendered with size 20 nm instead of the actual 5 nm for visualisation purposes and are coloured according to orientation (see wheel), scale bar is 200 nm. The video includes a timestamp for which $t = 0$ corresponds to the onset of the chemical pattern.

---

[7] Christian Vanhille-Campos and Šarić lab. Treadmilling filaments repository. `https://github.com/Saric-Group/treadmilling`.

[8] Jacob R. Gissinger, Benjamin D. Jensen, and Kristopher E. Wise. Modeling chemical reactions in classical molecular dynamics simulations. *Polymer*, 128:211–217, 2017.

[9] Jacob R. Gissinger, Benjamin D. Jensen, and Kristopher E. Wise. Reacter: A heuristic method for reactive molecular dynamics. *Macromolecules*, 53(22):9953–9961, 2020.

[10] Christian Vanhille-Campos and Šarić lab. Public repository storing the code and simulation data for this work. `https://doi.org/10.5522/04/24754527`.

[18] Natalia Baranova, Philipp Radler, Víctor M. Hernández-Rocamora, Carlos Alfonso, Mar López-Pelegrín, Germán Rivas, Waldemar Vollmer, and Martin Loose. Diffusion and capture permits dy-

namic coupling between treadmilling FtsZ filaments and cell division proteins. *Nature Microbiology*, 2020.

[19] Harold P Erickson, David E Anderson, and Masaki Osawa. FtsZ in bacterial cytokinesis: cytoskeleton and force generator all in one. *Microbiol Mol Biol Rev*, 74(4):504–28, 2010.

[20] Zsuzsanna Püspöki, Martin Storath, Daniel Sage, and Michael Unser. *Transforms and Operators for Directional Bioimage Analysis: A Survey*, pages 69–93. Springer International Publishing, Cham, 2016.

[21] Zuzana Dunajova, Batirtze Prats Mateu, Philipp Radler, Keesiang Lim, Dörte Brandis, Philipp Velicky, Johann Georg Danzl, Richard W Wong, Jens Elgeti, Edouard Hannezo, and Martin Loose. Chiral and nematic phases of flexible active filaments. *Nature Physics*, page 2022.12.15.520425, oct 2023.
